# The intestinal microbiota contributes to the development of immune-mediated cardiovascular inflammation and vasculitis in mice

**DOI:** 10.1101/2024.05.28.596258

**Authors:** Prasant K. Jena, Daiko Wakita, Angela C. Gomez, Thacyana T. Carvalho, Asli E. Atici, Meena Narayanan, Youngho Lee, Michael C. Fishbein, Patrice D. Cani, Willem M. de Vos, David M. Underhill, Suzanne Devkota, Shuang Chen, Kenichi Shimada, Timothy R. Crother, Moshe Arditi, Magali Noval Rivas

**Affiliations:** Department of Pediatrics, Division of Infectious Diseases and Immunology, Guerin Children’s at Cedars-Sinai Medical Center, Los Angeles, CA, USA; Infectious and Immunologic Diseases Research Center (IIDRC), Department of Biomedical Sciences, Cedars-Sinai Medical Center, Los Angeles, CA, USA; Department of Pathology and Laboratory Medicine, David Geffen School of Medicine, University of California Los Angeles, CA, USA; Louvain Drug Research Institute (LDRI), Metabolism and Nutrition Research Group (MNUT), UCLouvain, Université catholique de Louvain, Brussels, Belgium; WELBIO-Walloon Excellence in Life Sciences and BIOtechnology, WELBIO department, WEL Research Institute, Wavre, Belgium; Institute of Experimental and Clinical Research (IREC), UCLouvain, Université catholique de Louvain, Brussels, Belgium; Laboratory of Microbiology, Wageningen University, Wageningen, The Netherlands; Human Microbiome Research Program, Faculty of Medicine, University of Helsinki, Helsinki, Finland; Department of Biomedical Sciences, Cedars-Sinai Medical Center, Los Angeles, CA, USA; F. Widjaja Inflammatory Bowel Diseases Institute, Cedars-Sinai Medical Center, Los Angeles, CA, USA; Human Microbiome Research Institute, Cedars-Sinai Medical Center, Los Angeles, CA, USA; Smidt Heart Research Institute, Cedars Sinai Medical Center, Los Angeles, CA, USA

**Author notes:** Lead contact: Magali Noval Rivas, PhD, Cedars-Sinai Medical Center, Davis Research Building, 4th Floor, Suite D4018, 8700 Beverly Blvd. Los Angeles, California 90048, USA.

**Keywords:** Kawasaki disease, gut microbiota, vasculitis, coronary arteritis, *A. muciniphila*, *F. prausnitzii*, IL-1β, Amuc_1100

## Abstract

Alterations in the intestinal microbiota contribute to the pathogenesis of various cardiovascular disorders, but how they affect the development of Kawasaki disease (KD), an acute pediatric vasculitis, remains unclear. We report that depleting the gut microbiota reduces the development of cardiovascular inflammation in a murine model mimicking KD vasculitis. The development of cardiovascular lesions was associated with alterations in the intestinal microbiota composition and, notably, a decreased abundance of *Akkermansia muciniphila* and *Faecalibacterium prausnitzii.* Oral supplementation with either of these live or pasteurized individual bacteria, or with short-chain fatty acids (SCFAs) produced by them, attenuated cardiovascular inflammation. Treatment with Amuc_1100, the TLR-2 signaling outer membrane protein from *A. muciniphila*, also decreased the severity of vascular inflammation. This study reveals an underappreciated gut microbiota-cardiovascular inflammation axis in KD vasculitis pathogenesis and identifies specific intestinal commensals that regulate vasculitis in mice by producing metabolites or via extracellular proteins acting on gut barrier function.

**IN BRIEF:** It remains unclear whether changes in the intestinal microbiota composition are involved in the development of cardiovascular lesions associated with Kawasaki disease (KD), an immune-mediated vasculitis. Jena *et al.* observe alterations in the intestinal microbiota composition of mice developing vasculitis, characterized by reduced *A. muciniphila* and *F. prausnitzii*. Oral supplementation with either of these bacteria, live or pasteurized, or with bacteria-produced short-chain fatty acids (SCFAs) or Amuc_1100, the TLR-2 signaling outer membrane protein of *A. muciniphila*, was sufficient to alleviate the development of cardiovascular lesions in mice by promoting intestinal barrier function.

**HIGHLIGHTS:**

- Absence or depletion of the microbiota decreases the severity of vasculitis in a murine model mimicking KD vasculitis.
- Supplementation of *B. wadsworthia* and *B. fragilis* promotes murine KD vasculitis.
- Decreased abundances of *F. prausnitzii* and *A. muciniphila* are associated with the development of cardiovascular lesions in mice.
- Supplementation with either live or pasteurized *A. muciniphila* and *F. prausnitzii,* or the TLR-2 signaling Amuc_1100, reduces the severity of vasculitis by promoting gut barrier function.

## INTRODUCTION

Kawasaki disease (KD) is an acute febrile pediatric vasculitis that leads to coronary artery (CA) aneurysms in up to 25% of untreated children and is the primary cause of acquired heart disease in children in the United States ^1^. Timely treatment with intravenous immunoglobin (IVIG) significantly reduces the incidence of CA aneurysm development ^1^. However, up to 20% of KD patients are resistant to this treatment, have a higher risk of developing cardiac complications, and require additional therapies ^1^. Although the KD etiological agent(s) remains unidentified, the disease epidemiology, clinical presentation, and associated immune response indicate an infectious origin ^2^. KD is associated with systemic inflammation in several organs, including the lungs, liver, and gastrointestinal (GI) tract, and aneurysm development in medium-sized peripheral arteries ^1^. Infiltrations of innate immune cells have been reported in the inflamed CA of KD patients, which co-occur with elevated circulating levels of proinflammatory cytokines^3–6^.

While GI symptoms are not considered for KD diagnosis, abdominal pain, diarrhea, and vomiting are often reported at KD onset ^1,7^. These GI manifestations frequently complicate diagnosis, delay adequate treatment, and correlate with treatment resistance and the development of severe CA aneurysms ^7,8^. Notably, alterations in the intestinal microbiota composition have been reported during acute KD, often characterized by increased relative abundance of *Streptococcus* and *Staphylococcus* species and decreased abundance of bacteria producing short-chain fatty acids (SCFAs) ^9–14^. Furthermore, levels of the SCFA butyrate were found to be significantly lower in the stool of KD patients compared to controls ^15^. Perturbations of the intestinal microbiota composition and function have been associated with other inflammatory, autoimmune, metabolic, and cardiovascular disorders ^16–18^. However, the functional contributions of microbiota alterations to KD pathogenesis remain unknown.

Due to the overlapping clinical symptoms between KD and other pediatric infectious diseases, KD patients often receive antibiotic treatment pre-diagnosis ^19–21^. This treatment is typically ineffective and has even been associated with increased risk of IVIG resistance and higher rates of CA development ^2,19,22^. Antibiotics persistently impact the composition and function of the intestinal microbiota ^2,19,22,23^, thus these clinical findings support the hypothesis that microbiota composition may be a key determinant of KD pathogenesis.

Here, we examined the functional impact of the intestinal microbiota on KD using the *Lactobacillus casei* cell wall extract (LCWE)-induced KD vasculitis mouse model, which recapitulates key immunopathologic features of human KD, including coronary arteritis, aortitis, and luminal myofibroblast proliferation (LMP) in the CA lumen ^24–29^. We found that depletion of the gut microbiota in mice by antibiotic (Abx) treatment or absence of microbiota in germ-free (GF) mice attenuated the severity of LCWE-induced KD cardiovascular lesions. We observed alterations in the intestinal microbiota composition of mice after LCWE injection and evaluated the causal roles of these changes in promoting or decreasing the severity of vasculitis. We report reduced relative abundances of the SCFA-producing species *Akkermansia muciniphila* (*A. muciniphila*) and *Faecalibacterium prausnitzii* (*F. prausnitzii*), which correlate with the severity of LCWE-induced vasculitis. Replacing either live or pasteurized *A. muciniphila* or *F. prausnitzii*, or supplementing with acetate, propionate, butyrate, or Amuc_1100, a TLR-2 signaling protein present in the outer membrane of *A. muciniphila*, attenuated the severity of LCWE-induced KD cardiovascular inflammation by preventing LCWE-induced intestinal barrier dysfunction. Together, these results highlight a critical role of the gut microbiota in modulating cardiovascular inflammation in the LCWE model and suggest potential therapeutic strategies for KD.

## RESULTS

### The gut microbiota promotes LCWE-induced KD vasculitis

We used the LCWE-induced KD vasculitis mouse model to assess the potential contributions of the gut microbiota to vasculitis development. LCWE-injected specific pathogen-free (SPF) WT mice developed heart vessel inflammation characterized by aortitis, coronary arteritis, and abdominal aorta dilations (**Figure 1A, B**), as previously described ^24,26,27,29,30^. To gain insight into whether the gut microbiota promotes vascular inflammation in this model, we assessed the severity of LCWE-induced KD vasculitis in microbiota-depleted SPF mice treated with Abx (**Figure 1C, D,** and **Figure S1A**). SPF mice were provided an Abx cocktail in the drinking water either as a prophylactic treatment, given one week before LCWE injection or as a sustained Abx treatment, starting three days before LCWE injection until the experimental endpoint (**Figure S1A**). Both prophylactic and sustained Abx treatments resulted in a significant depletion of intestinal bacteria in the feces of treated animals (**Figure S1B**). Whereas LCWE-injected SPF mice developed severe heart vasculitis and abdominal aorta dilation and aneurysms, prophylactic and sustained Abx treatments were both associated with decreased severity of LCWE-induced heart vessel inflammation and development of abdominal aorta dilations (**Figure 1C, D**).

**Figure 1.**
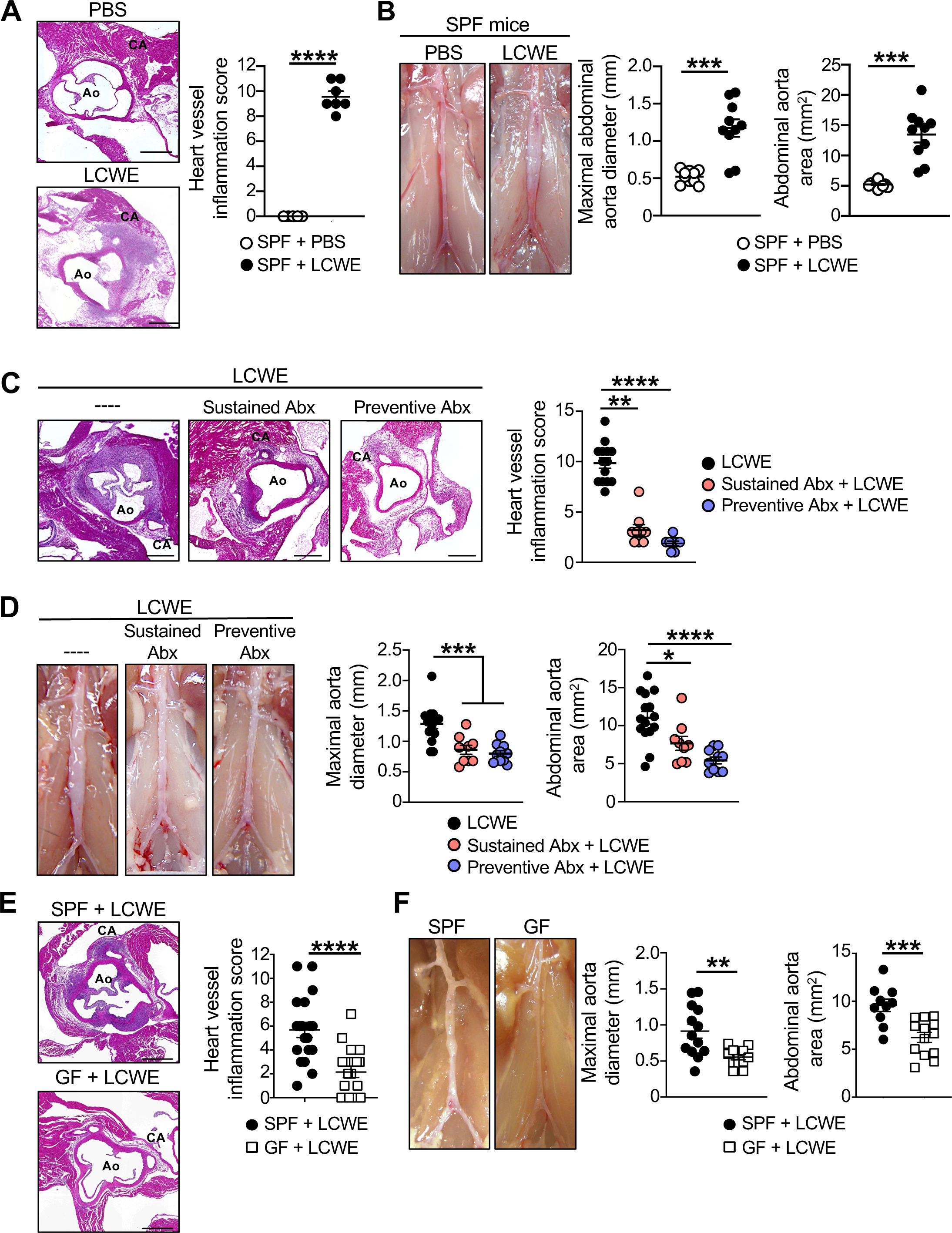
The intestinal microbiota promotes LCWE-induced KD vasculitis. **(A)** Representative H&E-stained heart sections and heart vessel inflammation score of SPF mice injected with either PBS or LCWE (n=5-7/group). Scale bars, 500µm. **(B)** Representative pictures of the abdominal aorta area, maximal abdominal aorta diameter, and abdominal aorta area measurements of SPF WT mice injected with either PBS or LCWE (n=10/group). **(C)** Representative H&E-stained heart sections and heart vessel inflammation score of LCWE-injected SPF mice and LCWE-injected SPF mice with preventative or sustained Abx treatment (n=8-14/group). Scale bars, 500µm. **(D)** Pictures of the abdominal aorta area, maximal abdominal aorta diameter, and abdominal aorta area measurements of LCWE-injected SPF mice and LCWE-injected SPF mice with preventative or sustained Abx treatment (n=9-15/group). **(E)** Representative H&E-stained heart sections and heart vessel inflammation score of LCWE-injected SPF and GF mice (n=19/group). Scale bars, 500µm. **(F)** Representative pictures of the abdominal aorta area, maximal abdominal aorta diameter, and abdominal aorta area measurements of SPF and GF mice injected with LCWE (n=10-13/group). Data are presented as mean ± SEM and representative of at least three independent experiments (A-B) or compiled from two-three independent experiments (C-F). \**p*<0.05, \*\**p*<0.01, \*\*\**p*<0.001, ****p<0.0001 obtained by Unpaired t-test with Welch’s correction (A, B, E, F), one way ANOVA with Kruskal-Wallis test with Dunn’s multiple comparisons (C), one-way ANOVA with Tukey’s multiple comparison tests (D). Abbreviations are as follows: Abx, antibiotics; CA, coronary artery; Ao, aorta; SPF, special pathogen-free; GF, germ-free.

Abx treatment during pregnancy limits microbiota diversity and its transfer from pregnant dams to their offspring. Therefore, in another set of experiments, we treated pregnant SPF dams with Abx, and their offspring were kept under the same treatment until LCWE injection at 5 weeks of age (**Figure S1C**). This resulted in depletion of intestinal bacteria in the feces of both Abx-treated pregnant dams and their offspring (**Figure S1D**). Furthermore, the offspring born from Abx-treated dams also exhibited decreased severity of LCWE-induced vasculitis, characterized by reduced heart inflammation and development of abdominal aorta dilations (**Figure S1E, F**). These results indicate that microbiota depletion decreases the severity of LCWE-induced KD vasculitis.

To further determine if mice reared in the absence of microbiota (germ-free, GF) were also protected, we next assessed the development of LCWE-induced cardiovascular inflammation in GF mice (**Figure 1E, F**). Indeed, the severity of cardiovascular lesions was significantly reduced in LCWE-injected GF mice compared to SPF mice (**Figure 1E, F**). Since mucosal and systemic immune responses are underdeveloped in GF mice ^31^, we next tested the effect of LCWE injection in GF mice raised with altered Schaedler flora (ASF). This well-defined bacterial consortium promotes typical gut structure and immune system development in mice ^32,33^. In this model, the severity of LCWE-induced KD vasculitis was significantly decreased compared to SPF mice (**Figure S1G, H**). These results reveal a crucial role of the intestinal microbiota in promoting the development of cardiovascular lesions in the LCWE-induced KD vasculitis in mice.

### LCWE-induced KD vasculitis is associated with alterations in gut microbiota composition

We next used 16S rRNA gene sequencing to determine changes in fecal microbiota composition associated with LCWE-induced KD vasculitis. Principal coordinate analysis (PCoA), as well as linear discriminant analysis effect size (LEfSe) analysis, indicated changes in the gut microbiota composition at 2 weeks post-LCWE injection when compared with baseline (day 0) (**Figures 2A and S2A**). LCWE-injected mice also showed increased alpha diversity and a higher number of observed operational taxonomic units (OTUs) than baseline mice, which may indicate some bacteria blooming (**Figure 2B**). Indeed, increased relative abundances of Firmicutes and Proteobacteria and decreased relative abundances of Actinobacteria and Verrucomicrobia were observed in the feces of LCWE-injected mice developing robust lesions (**Figure 2C**). To determine if either Firmicutes, mainly composed of Gram-positive bacteria, or Proteobacteria, mainly Gram-negative bacteria, contribute to LCWE-induced KD vasculitis, SPF mice received either Colistin or Vancomycin in the drinking water to deplete Gram-negative or Gram-positive bacteria, respectively (**Figure S2B, C**). Both Colistin and Vancomycin reduced the severity of LCWE-induced heart inflammation, indicating that Gram-positive and Gram-negative bacteria equally contribute to the development of LCWE-induced cardiovascular lesions (**Figure S2B, C**). The relative abundance of several bacterial families and genera was significantly altered in the LCWE-injected group (**Figures 2D-F and S2D**). Interestingly, we observed an increased relative abundance of members from the *Desulfovibrionaceae, Prevotellaceae,* and *Bacteroidaceae* families in LCWE-injected mice (**Figure 2E**). *Bilophila wadsworthia* (*B. wadsworthia*), which belongs to the Proteobacteria phylum and *Desulfovibrionaceae* family, and *Bacteroides fragilis* (*B. fragilis*), a member of the *Bacteroidaceae* family, are both opportunistic pathobionts capable of blooming in the intestinal tract and promoting intestinal inflammation and barrier dysfunctions ^34–36^. The relative abundance of bacteria belonging to the *Bacteroides*, *Desulfovibrio*, and *Prevotella* genera was also increased in LCWE-injected mice (**Figures 2E and S2D**). In contrast, the abundance of bacteria from the *Akkermansia* and *Faecalibacterium* genera was reduced (**Figures 2F and S2D**). Members of the families and genera that exhibited reduced relative abundance in LCWE-injected mice, such as *Akkermansia muciniphila* (*A. muciniphila; Akkermansia*) and *Faecalibacterium prausnitzii* (*F. prausnitzii; Faecalibacterium*) are known for their anti-inflammatory properties, specifically in the context of host metabolism and intestinal inflammation, respectively ^37–40^. Overall, these results indicate alterations in the intestinal microbiota composition of LCWE-injected mice, characterized by the enrichment of bacteria with pro-inflammatory capacities and the reduction of bacteria with anti-inflammatory effects.

**Figure 2.**
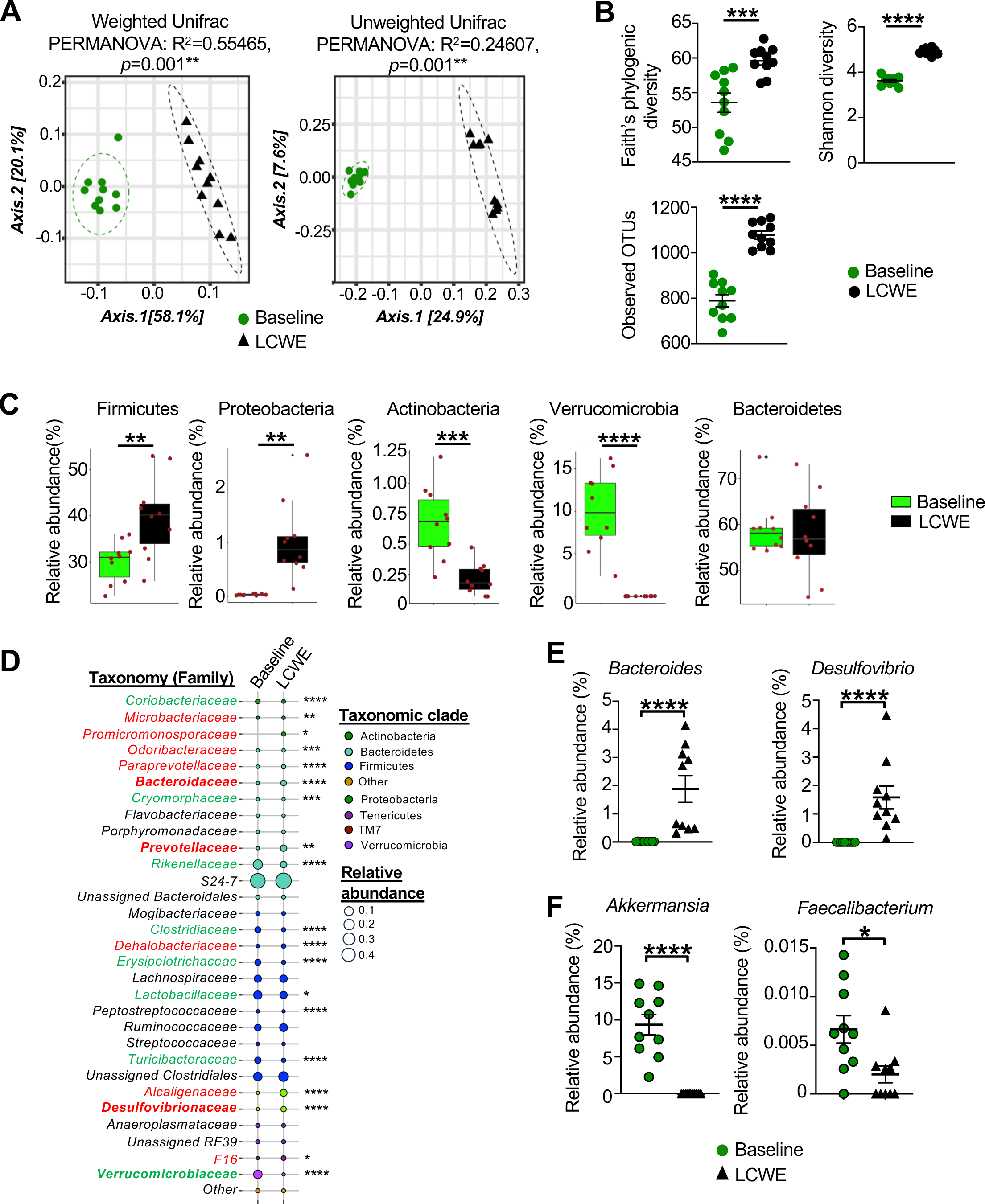
Intestinal microbiota composition changes associated with LCWE-induced KD vasculitis. **(A)** Principal coordinates analysis (PCoA) plots of weighted (left panel) and unweighted (right panel) UniFrac distance based on 16S rRNA gene sequencing of feces from SPF mice before (baseline) and 2 weeks after LCWE injection (n=10/group). **(B)** Alpha diversity index from 16S rRNA gene sequencing of feces from SPF mice before (baseline) and 2 weeks after LCWE injection. **(C)** Relative abundances of Firmicutes, Proteobacteria, Actinobacteria, and Verrucomicrobia in feces of SPF mice at baseline and 2 weeks after LCWE injection based on 16S sequencing results (n=10/group). **(D)** Bubble plot showing bacterial relative abundances at family level based on the 16S rRNA gene sequencing of feces of SPF mice at baseline and 2 weeks post-LCWE injection (n=10/group). Colors indicate the different taxonomic clades and bubble sizes of the relative abundance. **(E)** Relative abundances of genera *Bacteroides*, *Desulfovibrio*, *Akkermansia*, and *Faecalibacterium* in the fecal pellets from mice at baseline and 2 weeks after LCWE injection based on 16S sequencing results (n=10 mice/group). Data presented as mean +/- SEM, n=10 (A-E), \*\**p*<0.01, \*\*\**p*<0.001, \*\*\*\**p*<0.0001 by PERMANOVA (A), and Wilcoxon Rank Sum Test (B, C, E).

### B. wadsworthia and B. fragilis promote the severity of LCWE-induced KD vasculitis

We next confirmed the increased relative abundance of *B. wadsworthia* and *B. fragilis* by quantitative PCR (qPCR) analysis of feces collected from another independent cohort of PBS-control and LCWE-injected mice (**Figure S3A**). To determine if *B. wadsworthia* and *B. fragilis* functionally promote the development of LCWE-induced KD vasculitis, mice were treated with Abx for one week to deplete their microbiota and then orally administered every other day either a bacterial mix composed of *B. wadsworthia* and *B. fragilis*, starting one day before LCWE injection and until the experimental endpoint (**Figure S3B)**. Compared with LCWE-injected Abx-treated mice, relative abundances of *B. wadsworthia* and *B. fragilis* were increased in the feces of Abx-treated mice orally supplemented with the bacterial mix (**Figure 3A**). However, we noticed that 5 and 3 out of 19 LCWE-injected mice orally supplemented with the bacterial mix exhibited low levels of *B. wadsworthia* and *B. fragilis* enrichment (blue dots), respectively, similar to the levels detected in Abx-treated LCWE-injected mice (**Figure 3A**). Oral co-administration of *B. wadsworthia* and *B. fragilis* exacerbated LCWE-induced heart vessel inflammation compared to Abx-treated mice (**Figure 3B**), and tended to increase the development of abdominal aorta dilations and aneurysms (**Figure 3C, D**). Thus, reconstitution with these two species was sufficient to nearly normalize LCWE-induced vasculitis in Abx-treated mice.

**Figure 3.**
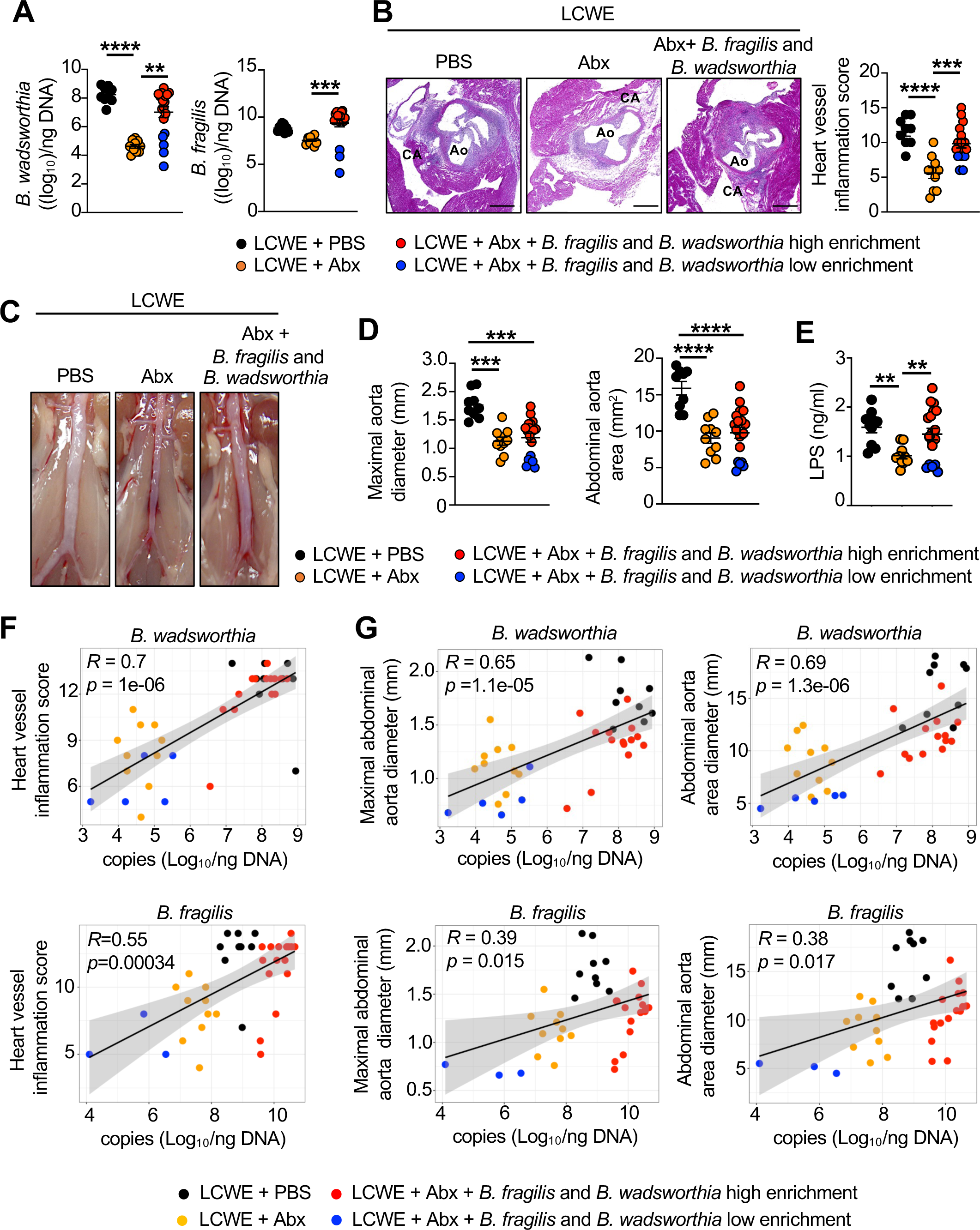
Oral supplementation with *B. wadsworthia* and *B. fragilis* exacerbates LCWE-induced KD vasculitis in Abx-treated SPF mice. **(A)** Quantification of *B. wadsworthia* and *B. fragilis* in fecal pellets from control mice that received PBS (n=9), Abx-treated mice (n=10), and Abx-treated mice orally supplemented with a bacterial mix composed of *B. wadsworthia* and *B. fragilis* (n=19) at 2 weeks-post LCWE injection. Among the Abx-treated mice supplemented with the bacterial mix, subgroups of high (red dots) or low (blue dot) enrichment of *B. wadsworthia* and *B. fragilis* were observed. **(B)** Representative H&E-stained heart sections and heart vessel inflammation score of control mice that received PBS (n=9), Abx-treated mice (n=10), and Abx-treated mice orally supplemented with *B. wadsworthia* and *B. fragilis* (n=17) at 2 weeks-post LCWE injection. Scale bars, 500µm. **(C, D)** Representative pictures of the abdominal aorta area (C), maximal abdominal aorta diameter, and abdominal aorta area measurements (D) of control mice that received PBS (n=9), Abx-treated mice (n=10), and Abx-treated orally supplemented with *B. wadsworthia* and *B. fragilis* (n=19) at 2 weeks-post LCWE injection. **(E**) Serum LPS concentrations of control mice that received PBS (n=10), Abx-treated mice (n=10), and Abx-treated orally supplemented with *B. wadsworthia* and *B. fragilis* (n=19) at 2 weeks-post LCWE injection. **(F)** Spearman correlations between heart vessel inflammation score and *B. wadsworthia* (top panel) and *B. fragilis* relative abundance in the fecal pellets of control mice (n=9), Abx-treated mice (n=10), and Abx-treated mice orally supplemented with *B. wadsworthia* and *B. fragilis* (n=19) at 2 weeks-post LCWE injection. **(G)** Pearson correlations between maximal abdominal aorta diameter and abdominal aorta area and *B. wadsworthia* and *B. fragilis* relative abundance in the fecal pellets of control mice that received PBS (n=9), Abx-treated mice (n=10), and Abx-treated mice orally supplemented with *B. wadsworthia* and *B. fragilis* (n=19) at 2 weeks-post LCWE injection. (A to G) Red and blue dots indicate mice SPF Abx-treated mice supplemented with the bacterial mix that exhibited high or low *B. wadsworthia* and *B. fragilis* enrichment, respectively. Data presented as mean ± SEM and pooled from two experiments \*\**p*<0.01, \*\*\**p*<0.001, \*\*\*\**p*<0.0001 by Kruskal-Wallis with Dunn’s multiple comparison tests (A, D left panel), one-way ANOVA with Tukey’s multiple comparisons tests (B, D right panel, E). Abbreviations are as follows: Abx, antibiotics; SPF, specific pathogen-free.

Changes in intestinal microbiota composition can alter gut permeability and increase lipopolysaccharide (LPS) dissemination to the systemic circulation, further promoting vascular inflammation, atherosclerosis, and thrombosis ^41^, and we have previously shown that LCWE-induced KD vasculitis is associated with a dysfunctional intestinal barrier and increased gut permeability ^26^. To determine how the intestinal microbiome affects LCWE-induced gut barrier function, we measured levels of circulating LPS. Abx-treated mice showed lower levels of circulating levels of LPS following LCWE injection than controls. In contrast, supplementation of Abx-treated mice with *B. wadsworthia* and *B. fragilis* increased the levels of circulating LPS (**Figure 3E**). Furthermore, the severity of LCWE-induced KD cardiovascular lesions, as measured by heart vessel inflammation, the maximal abdominal aorta diameter, and the abdominal aorta area, correlated strongly and positively with the relative abundance of *B. wadsworthia* and, to a lesser degree, *B. fragilis* (**Figures 3F, G**). Interestingly, we observed lower levels of circulating LPS and less severe LCWE-induced vascular inflammation in the subset of Abx-treated, bacteria-supplemented mice that exhibited low bacterial enrichment (**Figures 3B-F**). Overall, these results reveal that supplementation with *B. wadsworthia* and *B. fragilis* enhance the severity of LCWE-induced cardiovascular lesions.

### *F. prausnitzii* or *A. muciniphila* supplementation protects from LCWE-induced cardiovascular lesions

16S rRNA gene sequencing revealed decreased relative abundance of *Akkermansia* and *Faecalibacterium* genera in fecal pellets of LCWE-injected mice (**Figure 2F**). We next performed qPCR analysis of the feces from a novel and independent cohort of PBS control and LCWE-injected mice, and confirmed a significant reduction in *A. muciniphila* and *F. prausnitzii* levels in LCWE-injected mice (**Figure S4A**). Supplementation with live *F. prausnitzii* or *A. muciniphila* has been shown to benefit the host in several disorders ^40,42–44^. Furthermore, bacterial inactivation by pasteurization, a mild heat-mediated process, enhances their beneficial impact on the host, which may be mediated by increased stability and accessibility to bacterial proteins and components ^42,43^. To assess the functional implications of these two species on the development of LCWE-induced KD vasculitis, SPF mice were supplemented orally with either live or pasteurized *F. prausnitzii* daily, starting one week before LCWE injection until the experimental endpoint (**Figure 4A**). Mice supplemented with live *F. prausnitzii* had an increased relative abundance of *F. prausnitzii* in their feces (**Figure S4B**). Remarkably, supplementation with either live or pasteurized *F. prausnitzii* reduced the severity of LCWE-induced heart inflammation and the development of abdominal aorta dilations **(Figures 4B-D**). Similar results were obtained when supplementing mice with either live or pasteurized *A. muciniphila* (**Figures 4E-H, S4C**). Furthermore, the relative abundance of either *F. prausnitzii* or *A. muciniphila* in the feces of LCWE-injected mice supplemented with the live bacteria negatively correlated with heart vessel inflammation (**Figure S4D**) and the development of abdominal aorta aneurysms (**Figure S4E**). Next, we determined if pasteurized and inactivated *F. prausnitzii* or *A. muciniphila* reduced the severity of LCWE-induced KD vasculitis when provided in a therapeutic context, starting one day post-LCWE injection (**Figure 4I**). Treatment with either pasteurized *F. prausnitzii* or *A. muciniphila* significantly reduced the severity of heart vessel inflammation in LCWE-injected mice and tended to decrease the development of abdominal aorta dilations (**Figures 4G-I**). These findings reveal that supplementation with either live or pasteurized *F. prausnitzii* or *A. muciniphila,* both of which show reduced relative abundance during LCWE-induced KD vasculitis, has a beneficial effect and decreases the severity of LCWE-induced KD cardiovascular lesions development.

**Figure 4.**
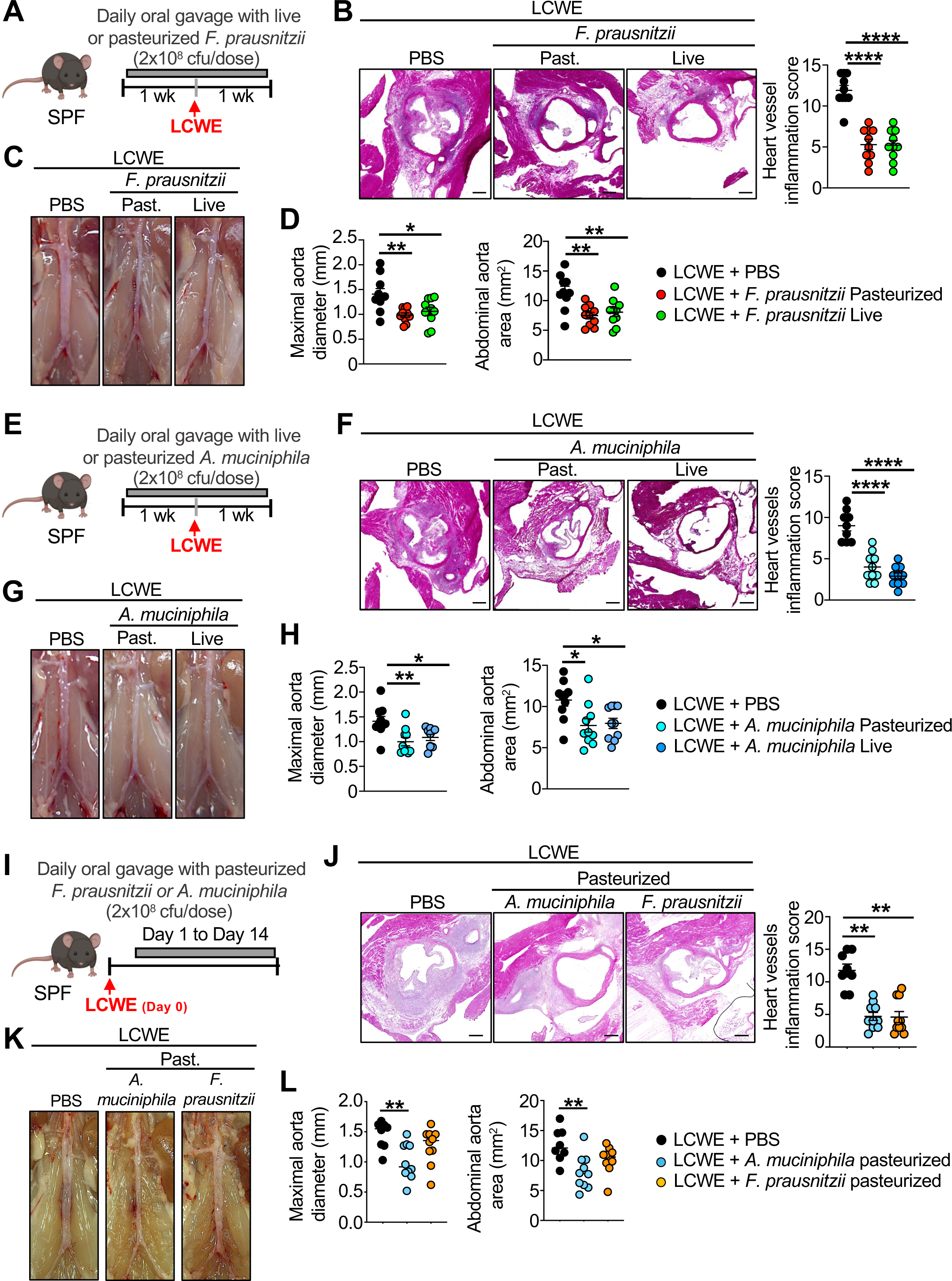
Live and pasteurized *F. prausnitzii* or *A. muciniphila* improve LCWE-induced KD vasculitis. **(A)** Schematic of the experimental design. Mice received by daily oral gavage live or pasteurized *F. prausnitzii* starting one week before LCWE injection and continuing until the experimental endpoint, at day 7 post-LCWE. **(B)** Representative H&E-stained heart sections and heart vessel inflammation score of LCWE-injected mice orally supplemented with either PBS, or live or pasteurized *F. prausnitzii* at one-week-post LCWE injection (n=10/group). Scale bars; 500µm. **(C, D)** Representative pictures of the abdominal aorta area (C), maximal abdominal aorta diameter, and abdominal aorta area measurements (D) of LCWE-injected mice untreated or orally supplemented with live or pasteurized *F. prausnitzii*, at one-week post-LCWE injection (n=10/group). **(E)** Schematic of the experimental design. Mice received by daily oral gavage live or pasteurized *A. muciniphila* starting one week before LCWE injection and continuing until the experimental endpoint, at day 7 post-LCWE. **(F)** Representative H&E-stained heart sections and heart vessel inflammation score of LCWE-injected mice orally supplemented with either PBS, or live or pasteurized *A. muciniphila,* at one-week-post LCWE injection (n=9-10/group). Scale bars; 500µm. **(G, H)** Representative pictures of the abdominal aorta area (G), maximal abdominal aorta diameter, and abdominal aorta area measurements (H) of LCWE-injected SPF mice untreated or orally supplemented with live or pasteurized *A. muciniphila* at one-week post LCWE injection (n=10/group). **(I)** Schematic of the experimental design. Mice were injected with LCWE and, one day later, were treated by oral gavage with either PBS, or pasteurized *F. prausnitzii* or *A. muciniphila* daily until day 14 post-LCWE injection. **(J)** Representative H&E-stained heart sections and heart vessel inflammation score of LCWE-injected mice untreated or orally supplemented with pasteurized *F. prausnitzii* or *A. muciniphila* (n=8-10/group). Scale bars; 500µm. **(K, L)** Representative pictures of the abdominal aorta area (K), maximal abdominal aorta diameter, and abdominal aorta area measurements (L) of LCWE-injected mice orally supplemented with either PBS, or pasteurized *F. prausnitzii*, or *A. muciniphila* (n=8-10/group). Data was compiled from two independent experiments (A to L) and presented as mean ± SEM. \**p*<0.05, \*\**p*<0.01, \*\*\*\**p*<0.0001 obtained by one-way ANOVA with Tukey’s multiple comparison tests (B, D, F, H right panel, L) or Kruskal-Wallis with Dunn’s multiple comparisons test (H left panel, J). Abbreviations are as follows: SPF, specific pathogen-free; cfu, colony forming unit.

### SCFAs decrease the severity of LCWE-induced KD vasculitis

The beneficial effects of *F. prausnitzii* and *A. muciniphila* could be mediated by secreted bacterial metabolites from the live bacteria, such as short-chain fatty acids (SCFAs), which are known for their anti-inflammatory properties and the enhancement of gut barrier function ^45^. To address this possibility, we next assessed the levels of SCFAs in the feces of PBS and LCWE-injected mice. We observed a significant reduction in acetate and propionate and a trend toward reduced butyrate fecal levels after LCWE injection (**Figure 5A**). Since *F. prausnitzii* uses acetate to generate butyrate ^46^ and *A. muciniphila* produces acetate and propionate ^37^, we tested if preventively supplementing these SFCAs in the drinking water would alter the development of LCWE-induced KD vasculitis (**Figure S5A**). Indeed, prophylactic supplementation with each SCFA significantly reduced the development of LCWE-induced heart inflammation and abdominal aorta dilations (**Figures 5B-D**). Compared with untreated LCWE-injected mice, SCFA supplementation also improved the gut barrier function, as demonstrated by reduced systemic levels of circulating LPS and D-lactate (**Figure 5E**). SCFA supplementation also increased the number of intestinal goblet cells and the expression of several intestinal tight junctions (TJs) in the gut tissues of LCWE-injected mice (**Figures 5F-H**). On the other hand, the expression of Claudin 2 (*Cldn2*), a TJ associated with a leaky intestinal barrier ^47^, was decreased by SCFAs supplementation (**Figure 5I**). Finally, SCFA supplementation increased the expression in intestinal tissues of the SCFA transporters MCT1, encoded by *Slc16a1*, and SMCT1, encoded by *Slc5a8* (**Figure 7I**).

**Figure 5.**
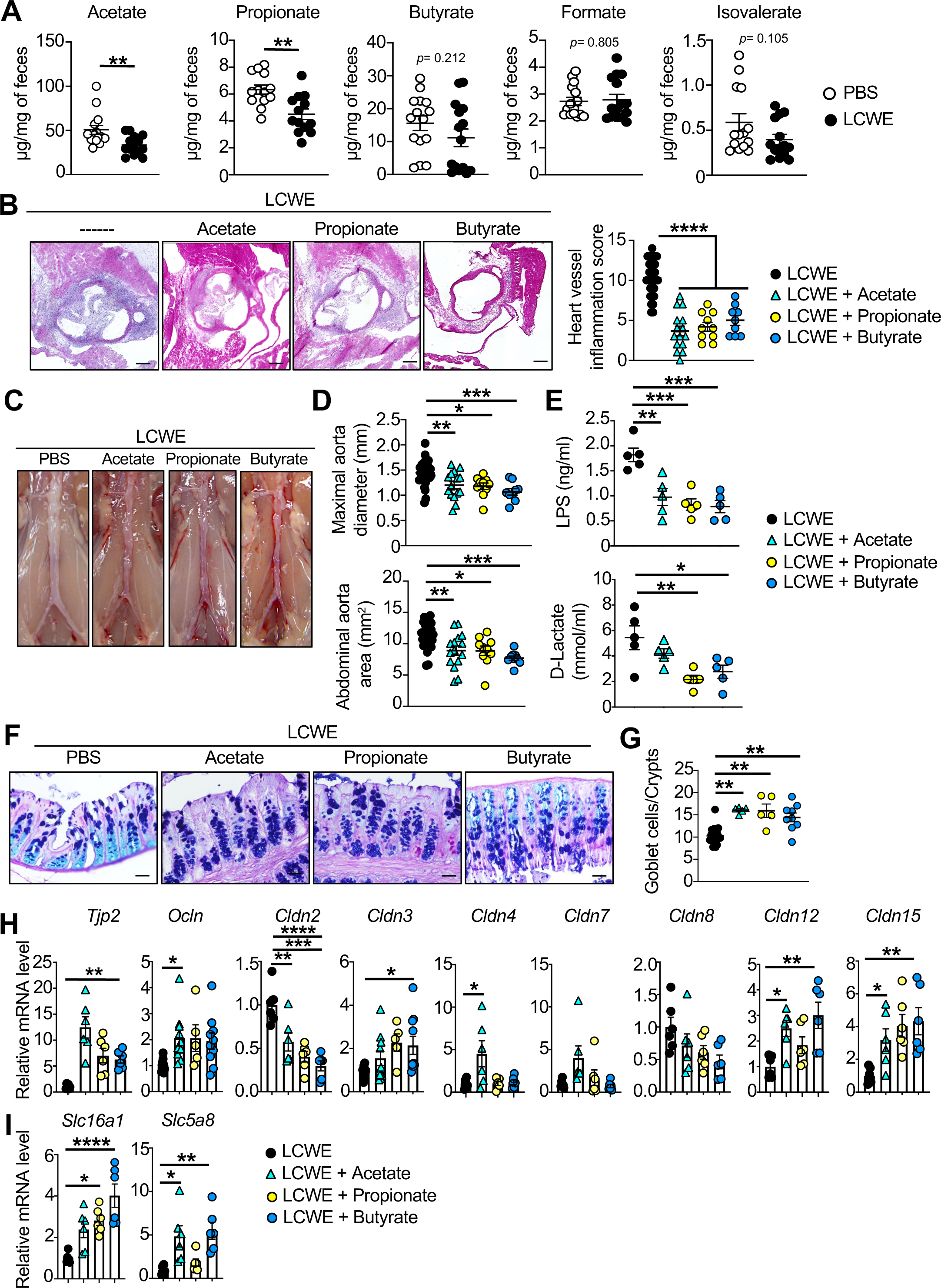
SCFAs oral supplementation reduces the severity of LCWE-induced KD vasculitis. **(A)** Quantification of acetate, propionate, butyrate, formate, and isovalerate in the fecal pellets of PBS and LCWE-injected mice at 2 weeks post-injection (n=15/group). **(B)** Representative H&E-stained heart sections and heart vessel inflammation score of untreated LCWE-injected mice and LCWE-injected mice orally supplemented with either acetate, propionate, or butyrate at one-week post-LCWE injection (n=9-32/group). Scale bars; 500µm. **(C, D)** Representative pictures of the abdominal aorta area (C), maximal abdominal aorta diameter, and abdominal aorta area measurements (D) score of untreated LCWE-injected mice and LCWE-injected mice orally supplemented with either acetate, propionate, or butyrate at one week-post LCWE injection (n=9-32/group)**. (E)** Serum LPS and D-Lactate concentrations of untreated LCWE-injected mice and LCWE-injected mice orally supplemented with either acetate, propionate, or butyrate at one week-post LCWE injection (n=9-32/group)**. (F, G)** Representative photomicrograph showing Alcian blue-periodic acid-Schiff (AB-PAS) staining of colon tissues (F) and Goblet cell counts per crypts of LCWE (n=13/group) from tissues of untreated LCWE-injected mice and LCWE-injected mice orally supplemented with either acetate, propionate, or butyrate at one weeks-post LCWE injection (n=5-9/group). Scale bars, 500µm. **(H, I)** mRNA levels of different TJs and SCFAs transporters in colon tissues of LCWE-injected mice (n=10) and LCWE-injected mice treated with sodium acetate (n=10), sodium propionate (n=5), and sodium butyrate (n=10) at one week post-LCWE injection. Data are presented as mean ± SEM and pooled from two-three independent experiments (A-G. \**p*<0.05, \*\**p*<0.01, \*\*\**p*<0.001 and \*\*\*\**p*< 0.0001 by unpaired t-test with Welch’s correction, two-tailed unpaired t-test (second from the left), one-way ANOVA with Tukey’s multiple comparison test (B, D, E, G, H right panel), or Kruskal-Wallis with Dunn’s multiple comparisons test (H left panel, I).

Since prophylactic supplementation with SCFAs reduced the severity of LCWE-induced KD vasculitis, we next asked if providing SCFAs therapeutically after LCWE injection would also impact the development of cardiovascular lesions. Mice were first injected with LCWE and supplemented with a mix of SCFAs in the drinking water on day 1, day 2, day 3, or day 5 post-LCWE injection (**Figure S5B**). We observed a significant reduction in heart inflammation and development of abdominal aorta dilations when SCFAs were provided in the drinking water on day 1 post-LCWE (**Figure S5C-E**). The therapeutic effect of SCFAs at day 1 post-LCWE injection was associated with increased intestinal mucus thickness and goblet cell numbers (**Figure S5F, G**). While delaying SCFA treatment to later time points tended to decrease heart vessel inflammation, this treatment did not reach significance or impact the development of abdominal aorta dilations (**Figure S5C-E**). These data indicate that supplementing SCFAs either prophylactically or therapeutically decreases the severity of LCWE-induced KD vasculitis and promote and restore intestinal barrier function.

### SCFAs and Amuc_1100 promote intestinal barrier function in an IL-1β dependent fashion

LCWE-induced KD vasculitis depends on NLRP3 activation and IL-1β production by monocytes and macrophages ^24,27–29^. Indeed, targeting the NLRP3-IL-1β axis using *Nlrp3^−/−^* and *Il1r1^−/−^* mice completely prevents the development of LCWE-induced KD vasculitis ^24,28,29^. Furthermore, LCWE induces an increase in intestinal permeability, which is IL-1β-mediated ^26^. Therefore, we next assessed *in vitro* the impact of SCFAs on the production of IL-1β by bone-marrow-derived macrophages (BMDMs) (**Supplementary Figure S6 A-D**). BMDMs were pretreated with either acetate, propionate, or butyrate for 8 hours and activated with either LPS and nigericin or LCWE (**Supplementary Figure S6 A-D**). While acetate and butyrate pretreatment of BMDMs decreased IL-1β production by both LPS/nigericin and LCWE-stimulated BMDMs, propionate only reduced IL-1β production by LCWE-stimulated BMDMs (**Supplementary** Figure 6B**, D**).

Caspase-1 is activated upon recruitment to an inflammasome and cleaves pro-IL-1β into its mature and biologically active form. We therefore assessed Caspase-1 activity by FAM-FLICA assay on heart tissues from LCWE-injected mice untreated or orally supplemented with the different SCFAs (**Supplementary** Figure 6E). SCFA supplementation reduced the numbers of macrophages with active caspase-1 (F4/80^+^ FLICA^+^) in heart tissue sections of LCWE-injected mice, with acetate having the most pronounced effect (**Supplementary** Figure 6E). We next examined the impact of oral supplementation of SCFAs on IL-1β production i*n vivo*. As previously published ^27^, LCWE injection resulted in increased levels of IL-1β in the peritoneal cavity at 24 hours post-injection, which were significantly reduced when the mice were orally administered a mix of SCFAs, composed of butyrate, propionate, and acetate (**Supplementary** Figure 6F). While we did not observe significant changes in levels of circulating LPS at 24 hours post-LCWE-injection, circulating concentrations of D-lactate were increased by LCWE and reduced by SCFA preventive supplementation (**Supplementary** Figure 6G). Overall, these results indicate that the SCFAs exert their beneficial effects during LCWE-induced KD vasculitis by promoting the intestinal barrier and targeting the IL-1β pathway.

Next, we evaluated the impact of exogenous IL-1β on intestinal organoid cultures generated from naïve WT or *Il1r1^−/−^* mice (**Figure 6A**). Cultured intestinal organoids generated from naïve WT mice and stimulated with recombinant IL-1β exhibited reduced organoid formation efficiency and surface area, decreased expression of intestinal TJs *Ocln* and *Cldn3,* and increased *Il1b* and *Il6* mRNA levels (**Figure 6A-D**). IL-1R specifically mediated these effects, as these responses were not observed in recombinant IL-1β-stimulated organoids generated from intestinal tissues of *Il1r1^−/−^*mice (**Figure 6A-D**).

**Figure 6.**
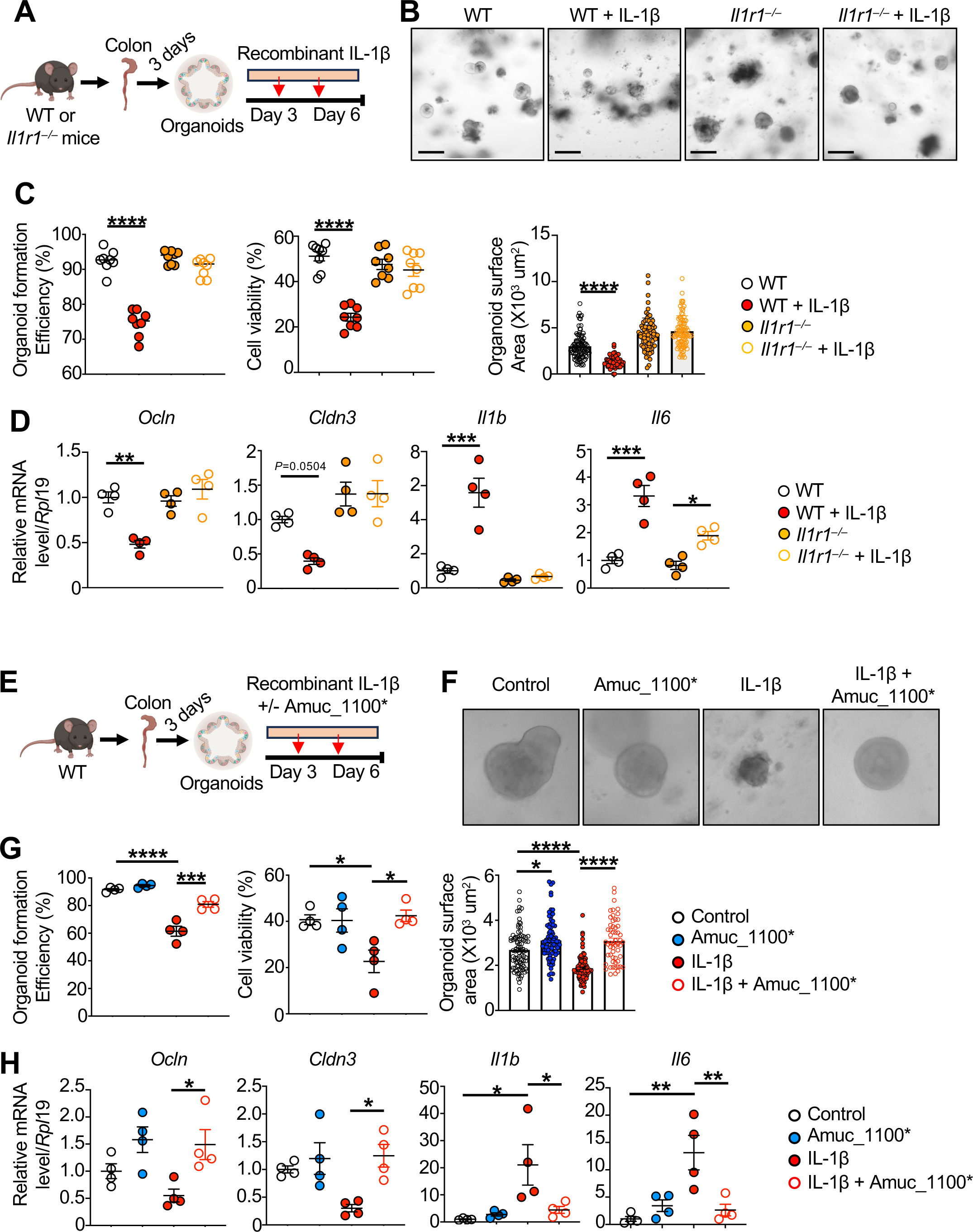
Amuc_1100* treatment counteracts IL-1β-mediated effects on intestinal organoids. **(A)** Schematic of the experimental design. Organoids were generated from the colons of naïve WT and *Il1r1^−/−^* mice and treated with recombinant IL-1β at the indicated days. **(B)** Representative pictures of organoids generated from naïve WT and *Il1r1^−/−^*mice, untreated or treated with IL-1β at day 3 and day 6 of the experiment. **(C)** Formation efficiency, viability, and surface area of organoids generated from naïve WT and *Il1r1^−/−^* mice, untreated or treated with IL-1β at day 3 and day 6 of the experiment (n=8/group). **(D)** quantified mRNA expression of intestinal TJs *Ocln* and *Cldn3,* and of *Il1b and Il6* in organoids generated from naïve WT and *Il1r1^−/−^* mice, untreated or treated with IL-1β at day 3 and day 6 of the experiment (n=4/group). **(E)** Schematic of the experimental design. Organoids were generated from the colons of naïve WT mice treated with recombinant IL-1β in the presence or absence of Amuc_1100*. **(F)** Representative pictures of organoids generated from naïve WT mice, treated with recombinant IL-1β in the presence of Amuc_1100*. **(G)** Formation efficiency, viability, and surface area of organoids generated from naïve WT mice, treated with recombinant IL-1β in the presence of Amuc_1100*(n=4/group). **(H)** quantified mRNA expression of intestinal TJs *Ocln* and *Cldn3,* and of *Il1b and Il6* in organoids generated from naïve WT mice, treated with recombinant IL-1β in the presence of Amuc_1100* (n=4/group). Data are presented as mean ± SEM and combined/pooled from two independent experiments. \**p*<0.05, \*\**p*<0.01, \*\*\**p*<0.001 and \*\*\*\**p*< 0.0001 by two-way ANOVA with Tukey’s multiple comparison test (C, D), one-way ANOVA with Tukey’s multiple comparison test (G left and middle panel, H) or Kruskal-Wallis with Dunn’s multiple comparisons test (G right panel).

As noted above, given that supplementation with either live or pasteurized *A. muciniphila* reduces the development of LCWE-induced cardiovascular lesions (**Figure 4E-L**), the effect of these bacteria may be mediated by secreted molecules, such as SCFAs, or by bacterial proteins that remain stable after pasteurization ^37,42,48^. Amuc_1100 is a 32 kDa protein abundantly expressed in the outer membrane of *A. muciniphila*, that signals to the TLR-2 receptor. A His-tagged Amuc_1100 (Amuc_1100*) produced in *E. coli* remains stable at 70°C, the temperature used for pasteurization, and recapitulates the beneficial effect of *A. muciniphila* on the gut barrier in the context of metabolic syndrome ^37,42^. We, therefore, evaluated the impact of Amuc_1100* on IL-1β-stimulated intestinal organoid cultures (**Figure 6E-I**). While stimulation with recombinant IL-1β decreased organoid growth, viability, the expression of intestinal TJs *Ocln* and *Cldn3*, and increased the expression of *Il1b* and *Il6*, Amuc_1100* addition reduced these IL-1β-mediated responses (**Figure 6F, H**). This suggests that Amuc_1100* is sufficient to promote intestinal barrier function by modulating intestinal TJ expression and reducing pro-inflammatory mediators, such as IL-1β and IL-6.

### Amuc_1100* attenuates LCWE-induced KD vasculitis

To further explore if Amuc_1100* may have a beneficial impact *in vivo*, we next compared the effects of supplementation with pasteurized *A. muciniphila* and Amuc_1100* on the development of LCWE-induced KD. WT mice received daily by oral gavage either PBS with glycerol, pasteurized *A. muciniphila,* or Amuc_1100*, starting one week before LCWE injection until the experimental endpoint (**Figure 7A**). Similar to the beneficial effect that we observed with pasteurized *A. muciniphila* (**Figure 4E-H**), Amuc_1100* led to a significant reduction in the severity of LCWE-induced KD vasculitis, as demonstrated by decreased heart vessel inflammation and reduced development of abdominal aorta dilations (**Figure 7B, C**). Compared with untreated LCWE-injected mice, which exhibit a reduced colonic mucus layer thickness, Amuc_1100* preserved the mucus thickness at levels similar to those observed in PBS control mice or LCWE-injected mice that received pasteurized *A. muciniphila* (**Figure 7D**). We next assessed the effect of Amuc_1100* on the expression of several TJ genes involved in regulating intestinal permeability. As expected, LCWE injection increased the mRNA expression of *Cldn2*, associated with increased gut permeability ^47^, and reduced the expression of *Tjp2*, *Ocln*, *Cldn3, Cldn12*, and *Cldn15*, but both Amuc_1100* and pasteurized *A. muciniphila* significantly prevented these effects *in vivo* (**Figure 7E**). Finally, treatment with both Amuc_1100* and pasteurized *A. muciniphila* restored the expression of the SCFA transporters MCT1, encoded by *Slc16a1*, and SMCT1, encoded by *Slc5a8* to levels similar to those observed in PBS control mice (**Figure 7F**). Altogether, these data further confirm that LCWE-induced KD vasculitis is associated with a dysfunctional intestinal barrier characterized by decreased expression of intestinal TJs, and that both Amuc_1100* and pasteurized *A. muciniphila* restore gut barrier function during LCWE-induced KD vasculitis and prevent the development of the cardiovascular lesions.

**Figure 7.**
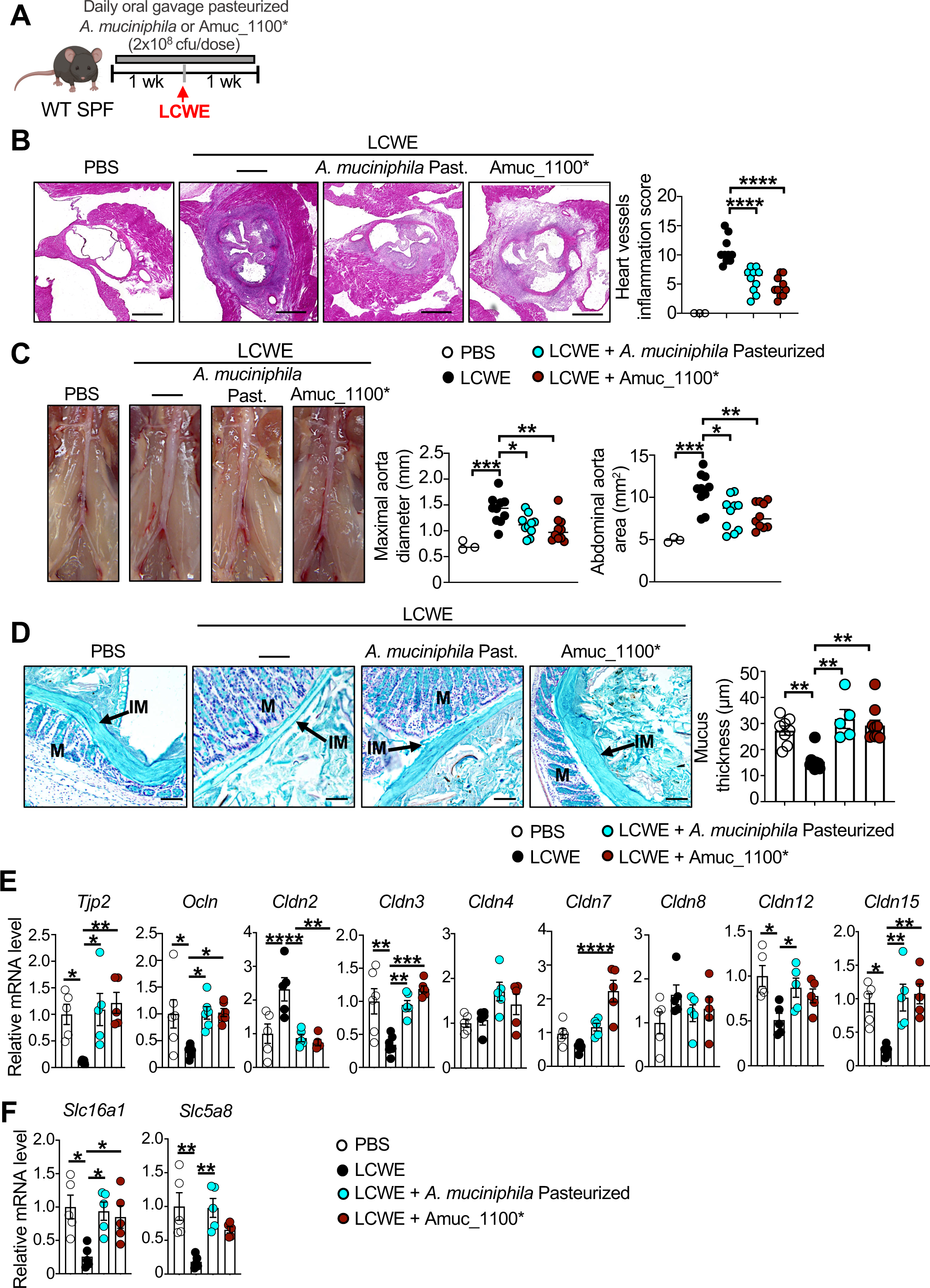
Pasteurized *A. muciniphila* and Amuc_1100 oral administration prevents LCWE-induced KD vasculitis. **(A)** Schematic of the experimental design. WT mice were supplemented with pasteurized *A. muciniphila* or Amuc_1100* daily by oral gavage starting 1 week before LCWE injection, and continuing until the end of the experiment at one-week post-LCWE. **(B)** Representative H&E-stained heart sections and heart vessel inflammation score of control mice and LCWE-injected mice orally supplemented with either PBS with 2.5% glycerol, or pasteurized *A. muciniphila* or Amuc_1100*\**, at one week-post LCWE injection (n=3-10/group). Scale bars; 500µm. **(C)** Representative pictures of the abdominal aorta area, maximal abdominal aorta diameter, and abdominal aorta area measurements of control mice and LCWE-injected mice orally supplemented with either PBS with 2.5% glycerol, or pasteurized *A. muciniphila* or Amuc_1100*\**, at one-week post-LCWE injection (n=3-10/group). **(D)** Representative Alcian-blue stained of colon tissue sections used to measure the mucus layer thickness of control PBS-injected mice and LCWE-injected mice orally supplemented with either PBS with 2.5% glycerol, or pasteurized *A. muciniphila* or Amuc_1100*, one-week post-LCWE injection (n=5-10/group). **(E, F)** Colon mRNA levels of intestinal TJs (E), and SCFA receptors *Slc16a1* and *Slc5a8* (F) in tissues from control PBS-injected mice and LCWE-injected mice orally supplemented with either PBS with 2.5% glycerol, or pasteurized *A. muciniphila* or Amuc_1100*, one-week post-LCWE injection (n=5-6/group). Data are presented as mean ± SEM and combined/pooled from two independent experiments *p<0.05, **p<0.01, ***p<0.001, and ****p< 0.0001 by one-way ANOVA with Tukey’s multiple comparison test. Abbreviations as follow: Past.; pasteurized, TJs; tight junction.

## DISCUSSION

The gut microbiota modulates homeostasis and immune responses in the GI tract and distant organs through the production of pathogen-associated molecular patterns and metabolites ^49^. Therefore, microbiota perturbations are often linked to immune dysfunction and disease development, including cardiovascular disorders ^17,18^. Here, we demonstrate several novel roles of the intestinal microbiota in the development of cardiovascular inflammation in a mouse model mimicking KD vasculitis. We show that the gut microbiota composition is altered in the setting of LCWE-induced vasculitis in mice. These changes in the intestinal microbiota composition could precipitate vasculitis development in mice or reflect a compensatory response of the gut microbiota to the intense host inflammatory response. Here, we demonstrate that these alterations contribute to the development of cardiovascular lesions and show that specific manipulations of the intestinal microbiota either promote or prevent the development of LCWE-induced vasculitis in mice. Furthermore, we demonstrate that *in vivo* supplementation with live or pasteurized *F. prausnitzii* or *A. muciniphila* decreases the severity of cardiovascular lesions in the LCWE model. This beneficial effect can be recapitulated by supplementing with SCFAs or with an outer membrane protein derived from *A. muciniphila*, Amuc_1100, that is known to signal to TLR-2, all of which promote intestinal barrier function and decrease the severity of vasculitis development.

KD patients often present with intestinal symptoms, such as vomiting, diarrhea, and abdominal pain ^7,8^. While the KD etiological agent(s) remains unidentified, it is suspected to be an infectious trigger(s) of viral origin with a mucosal portal of entry, possibly through airway epithelial cells or the intestinal barrier ^25,50^. KD affects mainly young children less than 5 years old, with the highest incidence in the first two years of life ^1,51^, but why KD develops in such an age-restricted population remains unknown. Given our demonstration of the key role of the microbiota in KD vasculitis, we speculate that it may be related to the fact that the early-life intestinal microbiota is less stable, exhibits a lower diversity and functional complexity and is more susceptible to environmental exposure, stabilizing into an adult-like conformation within the first three years of life ^52^. Interestingly, a retrospective study of an Italian cohort of KD patients indicated that younger patients (< 6 months) with abdominal and GI symptoms are at highest risk for developing CA aneurysms ^8^. Furthermore, immunohistochemical analysis of intestinal biopsies showed increased counts of HLA-DR^+^ cells and CD4^+^ T cells in the *lamina propria* from KD patients, indicating intestinal immune activation ^53^. These observations strongly point towards a “gut-cardiovascular” axis during KD vasculitis.

Here, we report that Abx-mediated acute depletion or complete lack of intestinal microbiota in GF mice significantly reduces the severity of LCWE-induced cardiovascular lesions, implying the contribution of the gut microbiota to vascular inflammation in this experimental model. Since KD manifests in children with prolonged fever, increased levels of inflammatory markers, and clinical features highly similar to an infectious disease, patients are often treated with Abx during the acute phase of the disease before the diagnosis of acute KD ^19,21,54^. Although intended to eliminate pathogenic organisms, this treatment dramatically impacts the gut microbiota composition and function ^55,56^, disrupting mucosal and immunological integrity, leading to the expansion of pathogenic bacteria and the loss of beneficial bacteria, thinning the intestinal mucus layer, and decreasing expression of intestinal TJs ^56,57^. In KD, Abx therapy it is associated with an increased rate of IVIG resistance ^19,22,50^. Therefore, it is possible that perturbations in the microbiota composition induced by Abx treatment provided pre-KD diagnosis may further promote the pro-inflammatory response associated with KD vasculitis and, potentially, IVIG resistance and even higher risk of development of coronary artery aneurysms ^2,19,22^.

Significant efforts have been made to determine if KD patients exhibit altered intestinal microbiota composition and to identify signature microbiota features associated with this disease ^9–14,58–60^. Initial culture-based studies from fecal samples and jejunal biopsies of KD patients indicated the loss of the *Lactobacillus* genus and the presence of Streptococci and Staphylococci ^9,14^. A metagenomic study analyzing feces from KD patients revealed the presence of strong alterations in the intestinal microbiota composition during the acute phase of the disease, and showed that the KD convalescence phase was associated with higher relative abundance of SCFA-producing bacteria ^11^. However, the reported alterations may reflect an Abx-mediated effect, as most enrolled patients were treated with Abx pre-KD diagnosis ^11^. A 16S rRNA gene sequencing-based longitudinal profiling of fecal microbiota communities of non-Abx-treated KD patients also demonstrated that the relative abundance of bacteria belonging to the *Enterococcus, Acinetobacter, Helicobacter, Lactococcus, Staphylococcus,* and *Butyricimonas* genera was increased during acute KD compared to age-matched healthy controls ^12^. Interestingly, several genera known to be SCFA producers were enriched in healthy controls and convalescent IVIG-treated KD patients when compared to acute KD patients ^12^. Furthermore, fecal butyrate levels were significantly lower in acute KD patients compared to normal controls ^15^. However, whether these alterations preceded KD or resulted from KD-induced inflammation remains unclear. Still, functional characterizations of the implicated bacterial genera affected are still lacking. Here, we report that oral supplementation with SCFAs decreases the severity of LCWE-induced KD vasculitis in mice, further hinting at a beneficial functional contribution of SCFAs-producing bacteria in this mouse model.

Here, we observed perturbations in the overall composition of fecal bacterial communities in mice developing LCWE-induced KD vasculitis, characterized by the blooming and fading of several bacterial families and genera. Specifically, we identified *B. wadsworthia (Desulfovibrionaceae)* and *B. fragilis (Bacteroidaceae)* as enriched in the feces of LCWE-injected mice and showed that oral supplementation with these two bacteria further promoted the severity of the disease. Both bacteria are opportunistic pathobionts capable of blooming in the intestinal tract, promoting intestinal inflammation and barrier dysfunction ^34–36^. *B. wadsworthia* is a sulfite-reducing pathobiont that expands in the presence of taurine-conjugated bile acids, a source of sulfur ^34^. Interestingly, liver panel test abnormalities and increased circulating levels of total bile acids have been reported in KD patients during the acute phase of the disease ^61,62^. In mice, *B. wadsworthia* is associated in mice with intestinal inflammation ^34^, atherosclerosis ^63^ and chronic metabolic conditions ^64^. Increased relative abundance of *B. wadsworthia* is also reported in the context of Behcet’s disease ^65^, a multisystem autoimmune disorder associated with whole-body vasculitis and juvenile idiopathic arthritis ^66^. Higher relative abundances of *Bacteroidaceae* have been previously reported in the fecal microbiota of patients with Henoch-Schönlein purpura, a vasculitis mediated by IgA immune complexes commonly affecting children from 4 to 7 years ^67,68^. The LCWE-induced KD vasculitis mouse model is also associated with IgA deposition in cardiovascular lesions and kidneys, a process that is IL-1β dependent ^26^. Indeed, IL-1β increases intestinal permeability ^69^, and a dysfunctional intestinal barrier may allow expanding luminal pathobionts and their metabolites to penetrate the *lamina propria* and trigger abnormal immune responses. We speculate that vascular diseases, such as IgA vasculitis and KD, may share a similar mechanistic pathway involving alterations in the intestinal microbiota composition and the blooming of pathobionts. However, the specific mechanisms and impact of expanding pathobionts on the host immune response will require further investigation.

In addition, we observed a decreased relative abundance of beneficial bacteria such as *F. prausnitzii* and *A. muciniphila* in the feces of LCWE-injected mice, which correlated with disease severity. The relative abundance of these two bacteria is often reported to be reduced in intestinal and extraintestinal disorders, including metabolic and cardiovascular diseases ^38,40,70^. Because of their robust positive systemic effects on host metabolism and anti-inflammatory properties, *Akkermansia* and *Faecalibacterium* are promising next-generation probiotics with a high preventive and therapeutic potential ^37,71^. We found that oral supplementation with *F. prausnitzii* or *A. muciniphila*, either prophylactically or therapeutically, significantly reduced LCWE-induced cardiovascular lesions. Mechanistically, this beneficial impact may be mediated in part by metabolites, such as butyrate and microbial anti-inflammatory molecule (MAM) ^72,73^ produced by *F. prausnitzii* ^46^, and acetate and propionate produced by *A. muciniphila* ^74^, which have anti-inflammatory and immune-modulatory properties and reinforce intestinal barriers ^71,73–75^. Indeed, we found that *in vivo* SCFA treatment prevented peritoneal production of IL-1β and decreased the number of infiltrating macrophages with Caspase-1 activity in inflamed heart tissues. Butyrate decreases the production of pro-inflammatory cytokines by suppressing NF-κb activation ^76^, and butyrate treatment of activated BMDMs reduces the secretion of nitric oxide, IL-6, and IL-12p40 by inhibiting histone deacetylases (HDAC) ^77^. Similarly, acetate and propionate also decrease IL-1β production ^78,79^. Here, we show that SCFA treatment also prevents IL1β production *in vitro* by LCWE-activated BMDMs. Furthermore, in mice, propionate was shown to have cardioprotective effect, and its administration attenuates vascular inflammation and atherosclerosis, an effect mainly mediated by regulatory T cells (Treg) ^80^. Here, we found that preventive and therapeutic oral supplementation with SCFAs significantly reduced the severity of LCWE-induced cardiovascular lesions. Altogether, these results suggest that diminished relative abundance of SCFA-producing microbiota in the gut of KD patients may play an important role in the development of cardiovascular lesions, but further investigations with larger number of patients are warranted.

We observed that providing *F. prausnitzii* or *A. muciniphila* in a pasteurized form reduced the severity of LCWE-induced KD vasculitis in mice to a similar level as supplementation with live bacteria or SCFAs. This is in line with previous studies demonstrating the beneficial effects of pasteurized *F. prausnitzii* and *A. muciniphila* in several disorders ^42,43,48,70^. Mice treated with pasteurized *A. muciniphila* exhibit increased goblet cell numbers and a thicker mucus layer ^42^, and we observed similar results in LCWE-injected mice. The fact that pasteurized *F. prausnitzii* and *A. muciniphila* exhibit a similar effect to live bacteria in preventing LCWE-induced KD vasculitis is exciting from a treatment perspective, as both bacteria are anaerobic and sensitive to oxygen, so their administration to patients in their live form may be challenging ^37^. Furthermore, the administration of live and pasteurized *A. muciniphila* is safe and tolerable in several proof-of-concept studies involving patients ^37,42,70^.

While pasteurization inactivates microorganisms and their production of SCFAs, this process also increases accessibility to extracellular bacterial proteins which resist heat denaturation ^42^. Amuc_1100 is an outer membrane protein specific to *A. muciniphila* which appear to remain stable at pasteurization temperature, and recapitulates the beneficial effect of live or pasteurized *A. muciniphila* in the context of metabolic disorders and colitis ^38,42,81–84^. Various studies have shown that purified Amuc_1100* is an efficient activator of the TLR-2 receptor ^37,42,85^. We show that Amuc_1100* also results in preventing LCWE-induced cardiovascular inflammation and vasculitis. Furthermore, using intestinal organoids and LCWE-injected mice, we show that while IL-1β decreases the expression of intestinal TJs, Amuc_1100* counteracts the effects of IL-1β and improves gut barrier function. Further studies are needed to determine if Amuc_1100* could potentially be used in patients preventively or therapeutically. In addition, further studies are required to identify which pasteurization-resistant *F. prausnitzii* proteins are involved in mediating its beneficial effect.

Since KD is the leading cause of acquired heart diseases in children in the USA, and up to 20% of KD patients are IVIG-refractory and at higher risk of developing cardiovascular lesions in childhood and long-term sequelae in adulthood, identifying novel intervention and therapeutic strategies is of critical interest ^86^. Overall, our study reveals a novel and important contribution of the gut microbiota in the development of cardiovascular lesions in a mouse model of KD vasculitis. Our results indicate that strategies aiming to modulate the production of microbial metabolites, or the microbiota composition, may be a preventive or adjunctive approach to current gold-standard therapy in treating cardiovascular disorders, including KD vasculitis. However, further studies are warranted to confirm our results in children with KD and to determine if preventive or therapeutic strategies modulating microbiota-host interactions and promoting gut barrier functions can safely and effectively be applied to KD patients.

## ACKNOWLEDGMENTS

We thank Malcolm Lane, Emily Aubuchon, and Debbie Moreira for their technical support, and laboratory members for their helpful discussions. We are grateful to Dr Anneleen Segers for her support in providing materials. M.N.R. is supported by NIH R01 HL139766 and R01 HL159297 grants. M.A. is supported by the NIH R01 AI072726 grant. M.A. and M.N.R. are supported by the NIH AI157274 grant. P.D.C. is an honorary research director at FRS-FNRS (Fonds de la Recherche Scientifique) and recipient of grants from FNRS (FRFS-WELBIO: WELBIO-CR-2022A-02 and FNRS-FWO: EOS: program no. 40007505).

## AUTHOR CONTRIBUTIONS

P.K.J., D.W., M.A., and M.N.R. conceived the project and designed the experiments. P.K.J, D.W., T.T.C., A.C.G., A.E.A., M.N. and M.N.R. performed experiments. M.C.F. provided a pathological review. D.U., S.D., S.C., T.R.C., K.S., and M.A. helped conceptualize, discuss results and provided critical reagents. W.M.d.V. and P.D.C. provided critical reagents, experimental design, and advice. P.K.J. and M.N.R. wrote the manuscript with input from all authors.

## DECLARATION OF INTERESTS

W.M.d.V. and P.D.C. are inventors on patent applications dealing with the use of specific bacteria and components in the treatment of different diseases. W.M.d.V. and P.D.C. have co-founded The Akkermansia Company SA, which is commercializing *Akkermansia muciniphila*, and have stocks in this start-up company. W.M.d.V. co-founded Caelus Health, and P.D.C. co-founded Enterosys. The remaining authors declare no competing interests.

## RESSOURCES AVAILABILITY

### Lead contact

Further information and requests for resources and reagents should be directed to and will be fulfilled by the Lead Contact, Magali Noval Rivas

### Materials availability

The study did not generate new, unique reagents.

### Data availability

16S rRNA gene sequencing data from this study are available from the Sequence Read Archive under the project ID “xxx”.

## EXPERIMENTAL MODELS AND SUBJECT DETAILS

### Mice

Specific pathogen-free (SPF) wild-type (WT) and *Il1r1^−/−^*C57BL/6J mice used in these experiments were purchased from Jackson Laboratories (Bar Harbor, ME, USA). Mice were housed under SPF conditions at Cedars-Sinai Medical Center and provided with a standard diet and water *ad libitum* unless noted otherwise for experimental purposes. Altered Schaedler Flora (ASF) mice were generated by colonizing germ-free (GF) mice with ASF stool. ASF and GF mice (Taconic) were kept under sterile conditions in flexible film isolators and exposed to a 14/10 h light/dark cycle. They were provided standard, autoclaved water, and gamma irradiated mouse chow *ad libitum*. Five week-old male animals were used in this study as LCWE injection induces more robust and more consistent coronary vasculitis lesions and abdominal aorta aneurysms in males compared with female mice ^29,87^. Animal experiments were conducted according to Cedars-Sinai Medical Center Institutional Animal Care and Use Committee guidelines.

### Bacteria

*Akkermansia muciniphila* (*A. muciniphila*; ATCC BAA-835) was grown anaerobically at 37**°**C (BD GasPak™ EZ container, Sparks, MD, USA) in Brain Heart Infusion (BHI) (Gibco, MD, USA) culture media supplemented with 0.5% mucin (porcine stomach, Type II, Sigma, USA) and 0.05% cysteine (Sigma-Aldrich) ^38,88^. *Faecalibacterium prausnitzii* (*F. prausnitzii;* ATCC 27768) was grown anaerobically at 37**°**C (BD GasPak™ EZ container, Sparks, MD, USA) in BHI culture media supplemented with 0.5% yeast extract (Fisher bioreagents, Cat#BP1422), cellobiose (1 mg/ml, Sigma Aldrich, USA), maltose (1 mg/ml, MP Biosciences), and cysteine (0.5 mg/ml, Sigma Aldrich) ^44,89^. *Bacteroides fragilis* and *Bilophila wadsworthia* cultures were a kind gift from Dr. Devkota (Cedars-Sinai Medical Center)^90^. *B. fragilis* was grown anaerobically at 37**°**C (BD GasPak™ EZ container, Sparks, MD, USA) in Brain Heart Infusion (BHI) and supplemented with hemin (5µg/ml, Sigma Aldrich, Cat#H9039) and vitamin K_1_ (1µg/ml; Sigma Aldrich, Cat#V3501). *B. wadsworthia* was grown anaerobically at 37**°**C (BD GasPak™ EZ container, Sparks, MD, USA) in dehydrated brucella media (Hardy diagnostics, CA, USA, Cat#C5311), supplemented hemin (5µg/ml, Sigma Aldrich, Cat#H9039) and vitamin K_1_ (1µg/ml; Sigma Aldrich, Cat#V3501), 0.5% sodium pyruvate (Sigma Aldrich, Cat#P-5280), and 0.5% taurine (Sigma Aldrich, Cat#T8691). Strains were confirmed by qPCR with specific primers. Bacterial concentrations were determined by serial dilution plating with the appropriate growth media and optical density measurement at 600nm.

## METHODS DETAILS

### The Lactobacillus casei cell wall extract (LCWE)-induced KD vasculitis murine model

LCWE was prepared as previously published ^24,91^. Five week old male mice were injected intraperitoneally (i.p.) with 500μl of LCWE or PBS (control group). One to two weeks after the LCWE injection, mice were euthanized, and blood was collected. Mice were then perfused with PBS, heart tissues were collected, and abdominal aortas were dissected as previously described ^29^. Abdominal aorta tissues were photographed before dissection and embedding in Tissue-Tek O.C.T. (Sakura Finetek). Maximal abdominal aorta diameter was determined by measuring 5 different areas separated by 2 mm of the abdominal aorta infra-renal portion (below the left renal artery) with ImageJ (NIH). The infrarenal abdominal aorta area was also measured in ImageJ. Abdominal aortas and heart tissues were embedded in O.C.T. and processed for histopathological examination.

### Tissue fixation, staining, and histological analysis

Serial cryosections (7μm) were prepared from the O.C.T.-embedded abdominal aorta and heart tissues and stained with hematoxylin and eosin (H&E, Sigma-Aldrich) for histological analysis. Histopathological examination and inflammation severity scoring of heart tissues were performed on H&E-stained sections by a senior investigator blinded to the experimental groups. Heart vessel inflammatory score (CAs, aortic root vasculitis, and myocarditis) was determined with a system previously described ^24,92^. Only heart sections showing the coronary artery branch separating from the aorta were scored. Briefly, acute inflammation, chronic inflammation, and connective tissue proliferation were each assessed using the following scoring system: 0 = no inflammation, 1 = rare inflammatory cells, 2 = scattered inflammatory cells, 3 = diffuse infiltrate of inflammatory cells, and 4 = dense clusters of inflammatory cells. Fibrosis was determined using the following scoring system: 0 = no medial fibrosis, 1 = medial fibrosis involving less than 10% of the CA circumference, 2 = medial fibrosis involving 11% to 50% of the CA circumference, 3 = medial fibrosis involving 51% to 75% of the CA circumference, and 4 = medial fibrosis involving more than 75% of the CA circumference. All 4 scores were combined to generate a severity score called “Heart inflammation score,” as previously published ^24,92^. Pictures were taken using a Biorevo BZ-9000 or BZ-X710 microscope (Keyence). Colon tissues were fixed in Carnoy’s solution (Medical Chemical) and embedded in paraffin blocks. Goblet cell numbers were assessed by Alcian blue-periodic acid Schiff’s (PAS) (Epredia) staining.

### Mucus Layer Thickness

Colon tissues were fixed in Carnoy’s solution (Electron Microscopy Sciences) for 12h at 4°C, followed by ethanol 100% for 24h, and embedded in paraffin. Tissue sections (7μm) were stained with Alcian Blue (Newcomer Supply) and analyzed either on a Biorevo BZ-9000 or BZ-X710 microscope (Keyence). A minimum of 10 measurements were made perpendicular to the inner mucus layer per field by using the ImageJ software (NIH).

### Antibiotic (Abx) treatment

Sterile drinking water supplemented with broad-spectrum antibiotics (Abx; ampicillin (1g/L), neomycin (1g/L), metronidazole (1g/L), vancomycin (500mg/L)) and sucralose (2gm/L) were provided *ad libitum* in the drinking water to SPF WT mice at the time-points specified in the experimental design. Control mice received normal sterile drinking water supplemented with sucralose (2gm/L). For preventive Abx treatment, Abx was provided to SPF WT mice starting one week before LCWE injection and removed on the day of LCWE injection. For sustained Abx treatment, Abx was provided three days before LCWE injection until the experimental endpoint. To deplete gram-positive or gram-negative bacteria, SPF WT mice received sterile drinking water supplemented with either vancomycin (500mg/L) or colistin (100mg/L), respectively, starting three days before LCWE injection until the experiment endpoint. Pregnant SPF WT dams were treated with Abx in the drinking water, starting two weeks before delivering their litter. For mock treatment, control dams received normal sterile drinking water. The litter of Abx-treated dams was kept on Abx treatment until 5 weeks, when they received normal sterile water, and the male pups were injected with LCWE.

### Bacteria pasteurization and supplementation to mice

2.0 x 10^8^ cfu of either *A. muciniphila* or *F. prausnitzii* in 200µl was inactivated by pasteurization for 30 min at 70°C. The absence of live bacteria was confirmed in pasteurized cultures by serial dilution plating with the appropriate growth media. Amuc_1100* is a His-tagged Amuc_1100 protein purified from overproducing *E.coli* produced as previously described ^42^ and provided by Dr. de Vos and Dr. Cani. Mice were treated with a daily oral administration of 5µg Amuc_1100* in 200µl of PBS containing 2.5% glycerol, starting one week before LCWE injection until the experiment endpoint. Control mice received 200µl of PBS containing 2.5% glycerol only. 2.0 x 10^8^ cfu combined of *B. wadsworthia* and *B. fragilis* in 200µl suspended in PBS was administered by oral gavage one dose before LCWE injection and every alternate day until the experiment endpoint.

### SCFA supplementation

SPF WT mice received *ad libitum* in the drinking water sodium butyrate (150mM, Sigma Aldrich), sodium acetate (200mM, Thermo Fisher Scientific), and sodium propionate (200mM, Sigma Aldrich), either separately or as a mix. Supplementation was started 3 days before LCWE injection and continued until the experiment endpoint. In another set experiment, the SCFAs mix (a cocktail of 150 mM sodium butyrate, 200mM sodium acetate, and 200mM sodium propionate) was provided starting at either day 1, day 2, day 3, or day 5 after LCWE injection.

### DNA extraction for bacterial sequencing

Fecal pellets of SPF mice were collected at the beginning of the experiment (baseline) and 2 weeks after LCWE injection. DNA for bacterial sequencing was extracted as previously described ^93,94^. Briefly, two fecal pellets (∼50mg) per sample were homogenized, incubated with lyticase (Sigma Aldrich, Cat#L-4025), bead beat (Omni Bead Rupture 12 Homogenizer, Omni International), and DNA was extracted with the QIAamp DNA mini kit (Qiagen, Cat#51304), according to the manufacturer’s instructions ^93,94^. DNA extracts were then submitted to the Cedars-Sinai Applied Genomics, Computation, and Translational (AGCT) Core for 16S rRNA sequencing.

### Library preparation, sequencing of bacterial amplicons and data analysis

Bacteria 16S rRNA gene amplicons were generated using high-fidelity Platinum™ SuperFi Polymerase (Life Technologies) using the primers 8F-overhang (5’-<u>TCGTCGGCAGCGTCAGATGTGTATAAGAGACAG</u>**AGAGTTTGATCMTGGCTCAG-**3’) and R357-overhang (5’-<u>GTCTCGTGGGCTCGGAGATGTGTATAAGAGACAG</u>**CTGCTGCCTYCCGTA-**3’) with the 48°C annealing temperatures and 25 PCR cycles. The underlined regions of the oligonucleotide sequence indicate the Illumina adapter sequence. The bolded region indicates the primer sequence. PCR reactions were purified using the ZR-96 DNA Clean-up Kit (Zymo Research). Amplicons were qualified on the 4200 TapeStation using the Agilent D1000 ScreenTape System (Agilent Technologies). Amplicons from up to 384 samples were uniquely indexed in a PCR reaction using Nextera XT Index Kit v2 Sets A-D (Illumina, San Diego, CA) and KAPA HiFi HotStart ReadyMix (Roche, Pleasanton, CA). Library enrichment was performed using 8 cycles of PCR. Multiplexed samples were pooled at equal volume by library type. The pooled libraries were purified using HighPrep PCR Magnetic Beads (MagBio Genomics) and subsequently assayed on a 4200 TapeStation to check final sizing. Samples were sequenced on the MiSeq platform (Illumina) with paired end 300bp sequencing chemistry. Raw data processing and run demultiplexing was performed using on-instrument analytics as per manufacturer recommendations. Raw FASTQ data were merged with overlap into single reads using SeqPrep v1.0 wrapped by QIIME v1.9 with default setting ^95^. A custom script was used to remove any reads that do not contain the proximal primer sequence or any reads containing a single N (unknown base) to enrich for high-quality reads. The remaining high-quality reads were aligned to the Greengenes reference database (May 2013 release) using BLAST v2.2.22 in QIIME v1.9 wrapper with an identity percentage ≥97% to select the operational taxonomic units (OTUs). All microbial community analyses were performed using R packages to assess 16S sequenced biom files by Phyloseq package, alpha, beta diversity, and permutational multivariate analyses of variance (PERMANOVA) by vegan package ^96^. Wilcoxon rank-sum tests were used to check the statistical significance among bacteria within each group using the wilcox.test function in R. The ggplot2 function was used to generate graphs.

### Bacterial Quantitative PCR

Quantifying *B. wadsworthia*, *A. muciniphila*, *B. fragilis*, and *F. prausnitzii* were performed by qPCR on DNA isolated from the fecal pellets of experimental mice using specific primers (**Key Resources Table**). Quantitative PCRs were performed with the PowerUp SYBR Green Master Mix (Applied Biosystems, Cat#A25741) on a CFX96 Real-Time system (Bio-Rad). qPCR mixtures contained 5µl of 2x PowerUp SYBR green Master Mix PCR conditions, 0.3µl of each forward and reverse primer (final concentration 10µM) and 2µl of DNA (40ng). PCR conditions were 95 °C for 10 minutes, followed by 40 cycles of 15 seconds at 95 °C, 30 seconds at 60 °C and, 32 seconds at 72 °C, 60 °C for 15 seconds. The bacterial gene copy number was determined using the Cp values and a standard curve generated by diluting the defined content of a pure sample of single bacterial species. All reactions were conducted in duplicate with appropriate negative controls.

### Fecal SCFA analysis

Fecal pellets from PBS and LCWE-injected mice were analyzed at the UC Davis West Coast Metabolomics Center for SCFAs quantification following standard operating procedures, as previously published ^97,98^.

### Quantitative Real-Time PCR analysis

Intestinal tissues were snap-frozen and homogenized in TRIzol (Invitrogen). RNA was extracted using the RNeasy kit (Qiagen). cDNA was prepared by reverse transcription using the RevertAid RT kit (Thermofisher, Cat#K1691). For reverse transcription, 100ng of RNA were transcribed into 20μl reaction volume containing 4µl 5X reverse transcription buffer, 2µl 10mM dNTP mix, 1µl RiboLock RNase Inhibitor (20U/µl), 1µl random hexamer primer, and 1µl reverse transcriptase. For reverse transcription, samples were mixed, centrifuged gently, and incubated for 5min at 25°C followed by 60min at 42°C using a thermal cycler (Applied Biosystem). qPCR analysis was performed the primers listed in the **Key Resource Table** and PowerUp SYBR Green Master Mix (Applied Biosystems, Cat#A25741) on a CFX96 Real-Time system (Bio-Rad). Relative expression of target genes was normalized to *Rpl19* ^81^. Samples were run in triplicate, and the relative expression was calculated using the ΔΔCt method.

### Serum ELISAs

Serum IL-1β was measured with the sensitive V-PLEX Mouse IL-1β Kit (Meso Scale Diagnostics, #K152QPD-1) per the manufacturer’s instructions. The samples were read and analyzed by MSD QuickPlex SQ120 instrumentation and Workbench 4.0 Software (Meso Scale Diagnostics). Serum LPS level was measured using the LPS ELISA kit (MyBiosource, MBS261904) and D-Lactate was measured using the Lactate acid assay kit (MyBiosource, MBS2567670)

### IL-1β quantification in murine peritoneal lavage

WT SPF mice were supplemented *ad libitum* in the drinking water with a mix of SCFAs (200mM acetate, 150mM butyrate, and 200mM propionate) for 3 days and injected with LCWE. The peritoneal lavage was collected 24 hours post-LCWE. Peritoneal lavage was centrifuged 3 minutes at 400g at 4°C and cells were discarded. IL-1β concentrations in the peritoneal lavage were assessed by ELISA (Thermofisher Scientific Cat# 887013A), according to the manufacturer’s instruction.

### Differentiation of bone marrow-derived macrophages (BMDMs)

Bilateral femora and tibiae of mice from WT mice were isolated to obtain bone marrow-derived macrophages (BMDMs). Bone marrow was flushed out of the bones with RPMI complete media (10% FBS and 1% PenStrep) and centrifuged at 400g for 3 min. The bone marrow cells were resuspended in RPMI complete media supplemented with 15% L-conditioned media and then placed into Petri dishes for macrophage differentiation. After 5–7 days of culture, BMDM (150,000 cells/well; 96 wells plate) were plated and incubated at 37°C for 24 hours. The cells were pretreated with 5mM of acetate, 5mM butyrate, and 5mM propionate for 8 hours. To activate the inflammasome, BMDMs were primed with 500 ng/ml of LPS for 4 hours and then stimulated with nigericin (10µm/ml) for 45 min. In another set of experiments, BMDM were stimulated with LCWE (10µg/ml) for 24 hours. Supernatants were collected to measure IL-1β level by ELISA (Thermofisher Scientific Cat# 887013A), according to the manufacturer’s instruction.

### Murine intestinal organoids

Colons were isolated, washed, and flushed with cold buffer (PBS with 1% penicillin and 1% streptomycin). Tissues were opened longitudinally, scrapped villi using cover slip and were cut into small pieces. Tissues were incubated on a rocker for 1 hour at 4°C in a chelation buffer (2mM EDTA (Hoefer, Cat #GR123), 5.6mM Na_2_HPO4 (Sigma, Cat #S0876), 8mM KH_2_PO4 (Sigma, Cat #P5379), 96.2mM NaCl (Sigma, Cat #S3014), 1.6mM KCl (Sigma, Cat #P9541), 43.4mM sucrose (Sigma, Cat #S0389), 54.9mM D-sorbitol (Sigma, Cat #S1876) and 12.5μl/50ml β-mercaptoethanol (Sigma, Cat #M6250)). Tissues were then filtered on 70 μm cell strainer, rinsed in cold PBS with 0.1% FBS (Gemini bioproducts, Cat #100106) and crypt enriched by centrifugation (1200 rpm, 3 min at 4°C). Isolated crypts were counted and embedded in Matrigel (Corning, cat #354263; 300 crypts/40μl) in a 24 wells plate in the presence of organoid growth medium (500µl of 1:1 mixture of L-WRN and DMEM/F12 with 5ml penicillin-streptomycin, 5ml glutamine (Sigma G7513), hygromycin (500 μg/mL), Gentamycin (500μg/mL) (Sigma, G81680), 5ml HEPES (10mM), 20% heat inactivated FBS, Murine EGF (50ng/ml) (Peprotech, Cat #31509), 10mM Y27632 dihydrochloride (Tocris, cat #1254). The organoid growth medium was changed every 3 days. Viability of colon organoids was monitored by Alamar Blue assays. To monitor viability at day 3 and day 9, we added a 1/10th volume of Alamar Blue reagent to the wells and incubated for 3h. Results were recorded using a fluorescence plate reader. After seeding the organoids, formation efficiency was counted manually by using a microscope on day 2 and at the end of the experiment. For formation efficiency, only viable, transparent, and well-formed organoids were considered, while dark, granulated, and immature organoids were excluded. To measure organoid surface area, the diameter of each organoid was measured in ImageJ. In a set of experiments, RNA was extracted from either intestinal crypts or organoid culture (6 days) generated from the colon of PBS and LCWE-injected mice. In another set, organoids generated from naïve mice were stimulated with IL-1β (10 ng/ml) and treated with Amuc_1100* (5ug /ml) on day 3 and day 6. On day 9, organoids were harvested, and RNA extracted with the RNeasy Micro kit (Qiagen, Cat#74004). Quantification of TJ genes was performed by qPCR with specific primers (**Key Resource Table**). Images of organoids were taken on day 3 and day 9 before harvest on a Biorevo BZ-9000 or BZ-X710 microscope (Keyence).

### Statistical analysis

Statistical analysis was performed using either R or Prism software v9 (GraphPad). The normality of data was assessed with the Shapiro-Wilk normality test with alpha=0.05. For comparisons of 2 groups, a 2-tailed unpaired Student’s t-test, with Welch’s correction when indicated, was used for normally distributed data. For nonparametric data, the Mann-Whitney two-tailed U/Wilcoxon rank test was used. For more than 2 group comparisons, 1-way ANOVA with Tukey post-test analysis was used for normally distributed data. The Kruskal-Wallis test with Dunn’s multiple comparisons test was used for non-normally distributed data. Statistical outliers were identified using the Grubbs’ test with a significance level of 0.05. Results are reported as means ± SEM, where each point represents one sample. A *p-value* of *p*<0.05 was considered statistically significant. No statistical methods were used to predetermine the sample size.

## SUPPLEMENTARY FIGURE LEGENDS

**Supplementary Figure 1 (*related to* Figure 1). Decreased severity of LCWE-induced KD vasculitis in both the offspring of Abx-treated dams and ASF-raised GF mice. (A)** Schematic of the experimental design. SPF mice received Abx in the drinking water either preventively starting one week before LCWE injection or sustainably throughout the course of the experiment, starting three days before LCWE injection until the experimental endpoint. **(B)** Quantitative PCR for bacterial 16S rDNA copies in the feces of SPF mice injected with LCWE that received a preventive or sustained Abx treatment (n=10/group). **(C)** Schematic of the experimental design. Pregnant SPF dams were treated with Abx in the drinking water. Their offspring were kept under Abx treatment until LCWE injection at 5 weeks of age. **(D)** Quantitative PCR for bacterial 16S rDNA copies in the feces of pregnant dams (left panel) and their offspring (right panel) treated or not with Abx in their drinking water (n=4-5/group). **(E)** Representative H&E-stained heart sections and heart vessel inflammation score of LCWE-injected SPF mice and Abx-treated offspring (n=11-16/group) at one-week post-injection. Scale bars, 500µm. **(F)** Representative pictures of the abdominal area, maximal abdominal aorta diameter, and abdominal aorta area measurements of LCWE-injected SPF mice and Abx-treated offspring at one-week post-injection and Abx-treated mice (n=10-13/group). **(G)** Representative H&E-stained heart sections and heart vessel inflammation score of PBS- and LCWE-injected SPF and GF mice raised with ASF (n=4-10/group) one-week post- LCWE injection. Scale bars, 500µm. **(H)** Representative pictures of the abdominal area, maximal abdominal aorta diameter, and abdominal aorta area measurements of PBS- and LCWE-injected SPF and GF mice raised with ASF (n=5-15/group) at one-week post-LCWE injection. Data presented as mean ± SEM and pooled from 2-3 independent experiments. *\*p*<0.05, *\*\*p*<0.01, \*\**\*p*<0.001, \*\*\**\*p*<0.0001 obtained by One-way ANOVA with Tukey’s multiple comparison tests (B, D), Unpaired t-test (D-F), two-way ANOVA with Tukey’s multiple comparison tests (G, H). Abbreviations are as follows: Abx; antibiotics, ASF; Altered Schaedler flora, GF, germ free.

**Supplementary Figure 2 (*related to*** Figure 2**). Alterations in gut microbiota composition associated to LCWE-induced KD vasculitis. (A)** Lefse analysis (cladogram) comparing bacterial communities by 16S rRNA gene sequencing of feces from SPF mice at baseline (day 0) and 2 weeks post-LCWE injection. (n=10/group). **(B)** Schematic of the experimental design. SPF mice received normal drinking water or drinking water supplemented with either Colistin or Vancomycin, starting 3 days before LCWE injection and until the experimental endpoint. **(C)** Representative H&E-stained heart sections and heart vessel inflammation score of LCWE-injected SPF mice untreated or treated with either Colistin or Vancomycin (n=4-5/group) at two-weeks post-injection. Scale bars, 500µm. **(D)** Heatmap showing relative abundances of bacteria at the genus level based on 16S rRNA gene profiling of the feces from SPF mice at baseline and 2 weeks after LCWE injection (n=10/group). **(E)** Red and green indicate increased or decreased relative abundance, respectively. Data presented as mean ± SEM. Data pooled from 2 independent experiments (E). \**p*<0.05, *\*\*p*<0.01, \*\**\*p*<0.001, and \*\*\**\*p*<0.0001 obtained by Kruskal-Wallis with Dunn’s multiple comparisons test (D, E), and Wilcoxon Rank Sum test (F).

**Supplementary Figure 3 (*related to*** Figure 3**): Relative abundances of *B. fragilis* and *B. wadsworthia* are increased after LCWE injection. (A)** Relative abundances of *B. fragilis* and *B. wadsworthia* rDNA copies measured by qPCR in the fecal pellets from an independent cohort of PBS and LCWE-injected mice at 2 weeks post-LCWE injection (n=10/group). **(B)** Schematic of the experimental design in which mice were treated with Abx in the drinking water for one week. After Abx water removal, mice were orally supplemented by oral gavage with a mix of *B. fragilis* and *B. wadsworthia* starting the day before LCWE injection, and then every other day. Data are presented as mean ± SEM and pooled from 2 independent experiments. \*\**p*<0.01 and \*\*\**\*p*<0.0001 by unpaired t-test (A).

**Supplementary Figure 4 (*related to*** Figure 4**). Decreased relative abundances of *F. prausnitzii* and *A. muciniphila* after LCWE injection. (A)** qPCR quantifications of the relative abundances of *F. prausnitzii* and *A. muciniphila* rDNA copies in the fecal pellets of PBS control mice or LCWE-injected mice at 2 weeks post-LCWE injection (n=10/group). **(B, C)** Relative abundances of *F. prausnitzii* (B) and *A. muciniphila* (C) rDNA copies in the fecal pellets of LCWE-injected mice orally supplemented with either live *F. prausnitzii* and *A. muciniphila* at 2 weeks post-LCWE injection (n=4-10/group). **(D, E)** Spearman or Pearson correlation between the relative abundance of *F. prausnitzii* and *A. muciniphila* and heart vessel inflation score (D) or maximal abdominal aorta diameter and total abdominal aorta area (Pearson correlation) (E). Data are presented as mean ± SEM and pooled from two independent experiments (A-C). \*\*\**p*<0.001 and \*\*\**\*p*<0.0001 by unpaired t-test with or without Welch’s correction (A), one-way ANOVA with Tukey’s multiple comparison tests (B), and Kruskal-Wallis with Dunn’s multiple comparisons test (C).

**Supplementary Figure 5 (*related to*** Figure 5**). Therapeutic SCFAs oral supplementation during LCWE-induced KD vasculitis. (A)** Schematic of the experimental design. Mice received *ad libitum* in drinking water either acetate, propionate, or butyrate starting 3 days before LCWE injection until the experiment endpoint. **(B)** Schematic of the experimental design. Mice received *ad libitum* in drinking water a cocktail of SCFAs composed of acetate, propionate and butyrate starting either on days 1, 2, 3 or 5 after LCWE injection and until the experiment endpoint. **(C)** Representative H&E-stained heart sections and heart vessel inflammation score of LCWE-injected mice and LCWE-injected supplemented with SCFAs starting at either day 1, 2, 3 or 5 post-LCWE injection (n=8-12/group) at two weeks post-LCWE. Scale bars, 500µm. **(D, E)** Representative pictures of the abdominal area (D), maximal abdominal aorta diameter and abdominal aorta area (E) of LCWE-injected mice and LCWE-injected supplemented with SCFAs starting at either day 1, 2, 3 or 5 post-LCWE injection (n=8-15/group) at two weeks post-LCWE **(F)** Representative photomicrograph showing Alcian blue staining of colon tissues and mucus thickness measurements from colon tissues of LCWE-injected mice and LCWE-injected mice supplemented with SCFAs starting at either day 1, 2, 3 or 5 post-LCWE injection (n=6-15/group) at two weeks post-LCWE. Scale bars; 100µm. **(G)** Representative photomicrograph showing Alcian blue-periodic acid-Schiff (AB-PAS) staining of colon tissues and Goblet cell counts per crypts of LCWE (n=10/group) from tissues from colon tissues of LCWE-injected mice and LCWE-injected supplemented with SCFAs starting at either day 1, 2, 3 or 5 post-LCWE injection (n=8-12/group) at two weeks post-LCWE. Scale bars, 500µm. Data was compiled from two-three independent experiments (C-G) and presented as mean ± SEM. \**p*<0.05, \*\**p*<0.01, \*\*\**p*<0.001, \*\*\*\**p*<0.0001 by Kruskal-Wallis with Dunn’s multiple comparisons test (C, E, F), and one-way ANOVA with Tukey’s multiple comparison test (G). Abbreviation of SCFA, short chain fatty acids.

**Supplementary Figure 6 (*related to*** Figure 5**). SCFAs decrease IL-1β production *in vitro* and *in vivo*. (A)** Schematic of the experimental design. BMDMs were pretreated with either acetate, propionate, or butyrate for 8 hours, primed with LPS, and stimulated with nigericin for 45m minutes, when supernatants were collected, and IL-1β was measured by ELISA. **(B)** Levels of IL-1β produced by BMDMs pretreated with either acetate, propionate, or butyrate, primed with LPS and stimulated with nigericin (n=3/group). **(C)** Schematic of the experimental design. BMDMs were pretreated with either acetate, propionate, or butyrate for 8 hours, and stimulated with LCWE for 24 hours, when supernatants were collected, and IL-1β was measured by ELISA. **(D)** Levels of IL-1β produced by BMDMs pretreated with either acetate, propionate, or butyrate, and stimulated with LCWE (n=4/group). **(E)** Representative picture of F4/80 and FLICA staining and quantification of F480+ FLICA+ cells in heart tissues from either PBS-injected mice, LCWE-injected mice, and LCWE-injected mice that were orally supplemented with either acetate, butyrate, or propionate at one week post-LCWE injection. SCFAs supplementation was started 3 days before LCWE injection. White arrows indicate F4/80+ FLICA+ cells (n =3-8/group). Scale bars: 50µm. **(F)** IL-1β levels in the peritoneal lavage of PBS-injected control mice, LCWE-injected mice, and LCWE-injected mice orally supplemented with a mix of SCFAs (butyrate, acetate, and propionate, provided 3 days before LCWE injection), at 24 hours post-LCWE injection (n=5-14/group). **(G)** Serum levels of LPS and D-Lactate from PBS-injected control mice, LCWE-injected mice, and LCWE-injected mice orally supplemented with a mix of SCFAs (butyrate, acetate, and propionate, provided 3 days before LCWE injection), at 24 hours post-LCWE injection (n=4-5/group). Data was compiled from two independent experiments (B-F) and presented as mean ± SEM. \**p*<0.05, \*\**p*<0.01, \*\*\**p*<0.001, \*\*\*\**p*<0.0001 by Kruskal-Wallis with Dunn’s multiple comparisons test (C, E) and one-way ANOVA with Tukey’s multiple comparisons (F, G).

**Supplementary Figure 7 (*related to*** Figure 7**). Decreased expression of *Cldn3* and *Ocln* mRNA by intestinal organoids from LCWE-injected mice. (A)** Measurements of colon lengths from PBS-injected control WT mice, and LCWE-injected WT and *Il1r1^−/−^* mice, at 2 weeks post-LCWE injection (n=8-10/group)**. (B)** Quantification of *Cldn3, Ocln, Il1b,* and *Il6* mRNA in intestinal crypts isolated from the colon of PBS and LCWE-injected mice (n=4-5/group)**. (C)** Schematic of the experimental design. Cell suspensions were generated from the colons of PBS and LCWE-injected WT mice, intestinal crypts and organoids were generated for *in vitro* studies. **(D)** Representative pictures of organoids generated from the colon of PBS and LCWE-injected mice (n=8-10/group). **(E)** Formation potential, viability, and surface area of organoids generated from the colon of PBS and LCWE-injected mice (n=8-10/group). **(F)** Quantification of *Cldn3, Ocln, Il1b,* and *Il6* mRNA in organoids generated from the colon of PBS and LCWE-injected mice (n=4-5/group). Data are presented as mean ± SEM and combined/pooled from two independent experiments. \**p*<0.05, \*\**p*<0.01 by Kruskal-Wallis with Dunn’s multiple comparisons test (A), two-tailed unpaired t-test (B, F left three panels), and unpaired t-test with Welch’s correction (F left panel).

