## supplementary file for "The intestinal microbiota contributes to the development of immune-mediated cardiovascular inflammation and vasculitis in mice"

### Figure S1 *(related to Figure 1)*

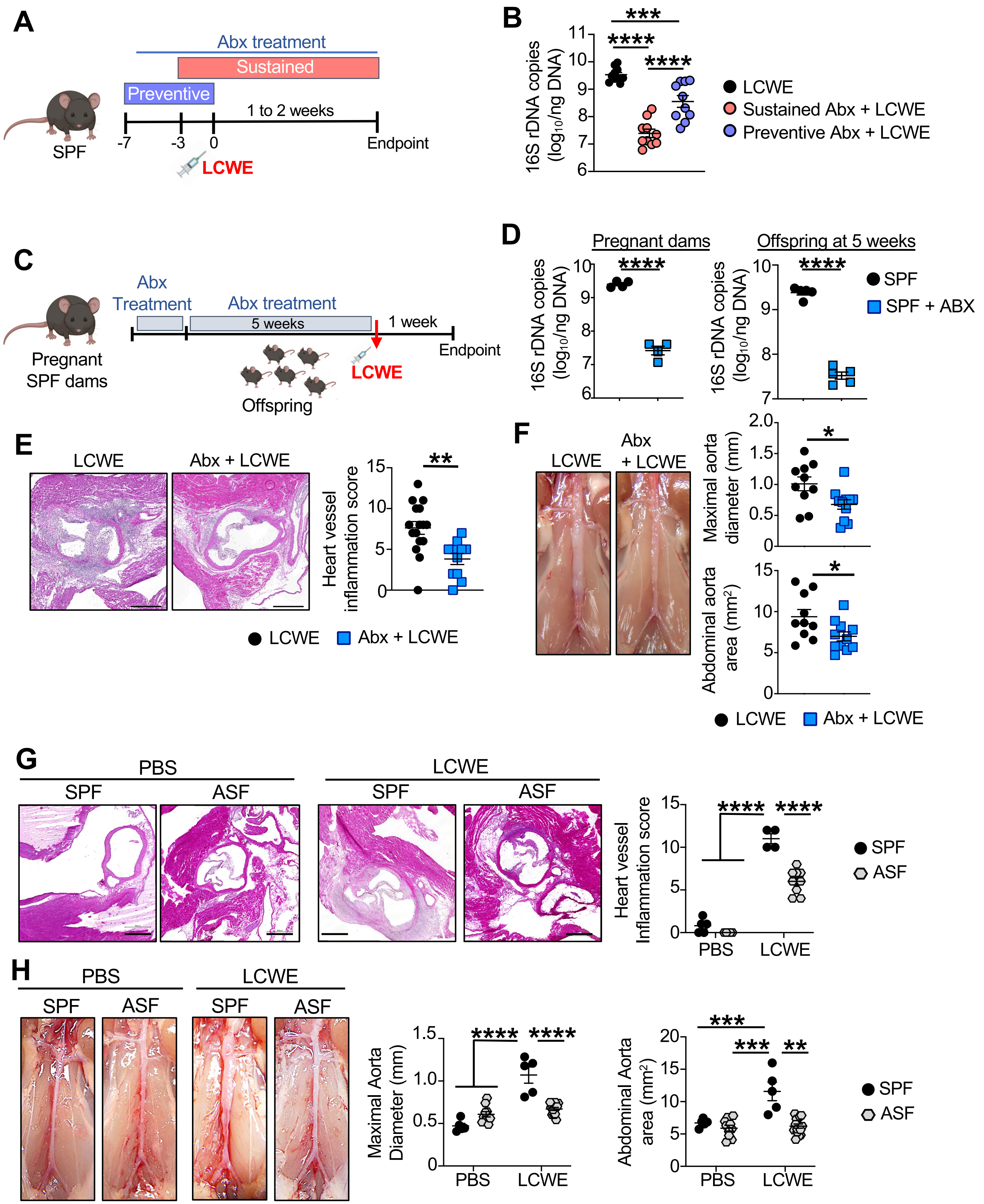

**Supplementary Figure 1 (*related to Figure 1*). Decreased severity of LCWE-induced KD vasculitis in both the offspring of Abx-treated dams and ASF-raised GF mice.**

**(A)** Schematic of the experimental design. SPF mice received Abx in the drinking water either preventively starting one week before LCWE injection or sustainably throughout the course of the experiment, starting three days before LCWE injection until the experimental endpoint. **(B)** Quantitative PCR for bacterial 16S rDNA copies in the feces of SPF mice injected with LCWE that received a preventive or sustained Abx treatment (n=10/group). **(C)** Schematic of the experimental design. Pregnant SPF dams were treated with Abx in the drinking water. Their offspring were kept under Abx treatment until LCWE injection at 5 weeks of age. **(D)** Quantitative PCR for bacterial 16S rDNA copies in the feces of pregnant dams (left panel) and their offspring (right panel) treated or not with Abx in their drinking water (n=4-5/group). **(E)** Representative H&E-stained heart sections and heart vessel inflammation score of LCWE-injected SPF mice and Abx-treated offspring (n=11-16/group) at one-week post-injection. Scale bars, 500µm. **(F)** Representative pictures of the abdominal area, maximal abdominal aorta diameter, and abdominal aorta area measurements of LCWE-injected SPF mice and Abx-treated offspring at one-week post-injection and Abx-treated mice (n=10-13/group). **(G)** Representative H&E-stained heart sections and heart vessel inflammation score of PBS- and LCWE-injected SPF and GF mice raised with ASF (n=4-10/group) one-week post-LCWE injection. Scale bars, 500µm. **(H)** Representative pictures of the abdominal area, maximal abdominal aorta diameter, and abdominal aorta area measurements of PBS- and LCWE-injected SPF and GF mice raised with ASF (n=5-15/group) at one-week post-LCWE injection. Data presented as mean ± SEM and pooled from 2-3 independent experiments. **p*<0.05, ***p*<0.01, ****p*<0.001, *****p*<0.0001 obtained by One-way ANOVA with Tukey's multiple comparison tests (B, D), Unpaired t-test (D-F), two-way ANOVA with Tukey's multiple comparison tests (G, H). Abbreviations are as follows: Abx; antibiotics, ASF; Altered Schaedler flora, GF, germ free.

### Figure S2 *(related to Figure 2)*

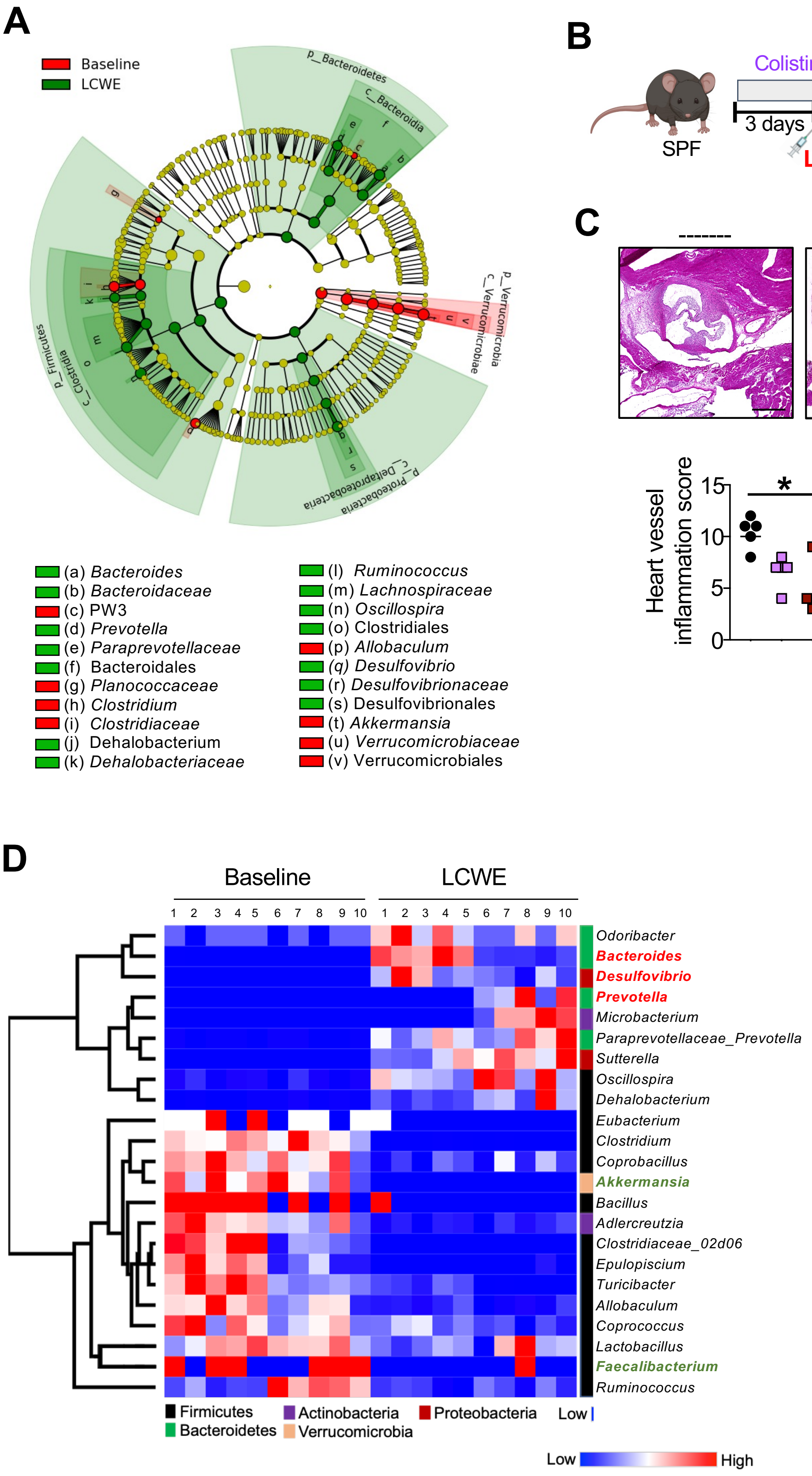

**Supplementary Figure 2 (*related to Figure 2*). Alterations in gut microbiota composition associated to LCWE-induced KD vasculitis*.***

**(A)** Lefse analysis (cladogram) comparing bacterial communities by 16S rRNA gene sequencing of feces from SPF mice at baseline (day 0) and 2 weeks post-LCWE injection. (n=10/group). **(B)** Schematic of the experimental design. SPF mice received normal drinking water or drinking water supplemented with either Colistin or Vancomycin, starting 3 days before LCWE injection and until the experimental endpoint. **(C)** Representative H&E-stained heart sections and heart vessel inflammation score of LCWE-injected SPF mice untreated or treated with either Colistin or Vancomycin (n=4-5/group) at two-weeks post-injection. Scale bars, 500μm. **(D)** Heatmap showing relative abundances of bacteria at the genus level based on 16S rRNA gene profiling of the feces from SPF mice at baseline and 2 weeks after LCWE injection (n=10/group). **(E)** Red and green indicate increased or decreased relative abundance, respectively. Data presented as mean ± SEM. Data pooled from 2 independent experiments (E). **p*<0.05, ***p*<0.01, ****p*<0.001, and *****p*<0.0001 obtained by Kruskal-Wallis with Dunn's multiple comparisons test (D, E), and Wilcoxon Rank Sum test (F).

### Figure S3 *(related to Figures 3)*

**A**

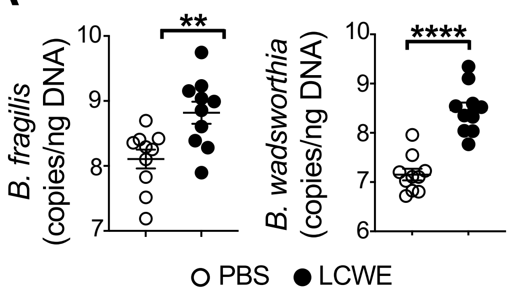


**B**

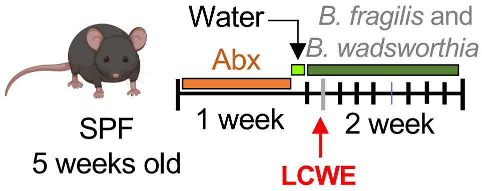


**Supplementary Figure 3 (*related to Figure 3*): Relative abundances of *B. fragilis* and *B. wadsworthia* are increased after LCWE injection.**

**(A)** Relative bundances of *B. fragilis* and *B. wadsworthia* rDNA copies measured by qPCR in the fecal pellets from an independent cohort of PBS and LCWE-injected mice at 2 weeks post-LCWE injection (n=10/group). **(B)** Schematic of the experimental design in which mice were treated with Abx in the drinking water for one week. After Abx water removal, mice were orally supplemented by oral gavage with a mix of *B. fragilis* and *B. wadsworthia* starting the day before LCWE injection, and then every other day. Data are presented as mean ± SEM and pooled from 2 independent experiments. ***p*<0.01 and *****p*<0.0001 by unpaired t-test (A).

### Figure S4 *(related to Figure 4)*

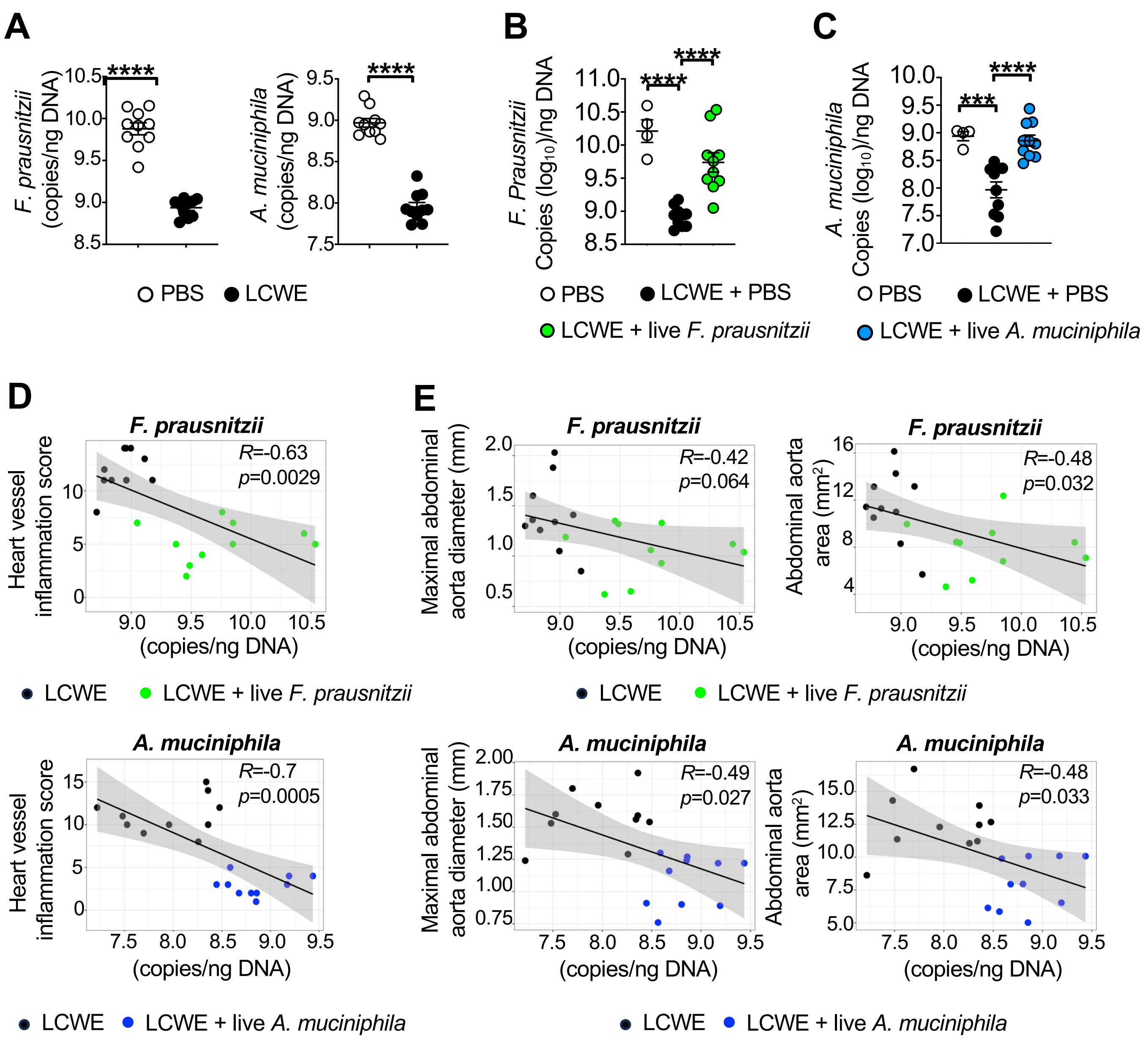


**Supplementary Figure 4 (*related to Figure 4*). Decreased relative abundances of *F. prausnitzii* and *A. muciniphila* after LCWE injection.**
**(A)** qPCR quantifications of the abundances of *F. prausnitzii* and *A. muciniphila* rDNA copies in the fecal pellets of PBS control mice or LCWE-injected mice at 2 weeks post-LCWE injection (n=10/group). **(B, C)** Abundances of *F. prausnitzii* (B) and *A. muciniphila* (C) rDNA copies in the fecal pellets of LCWE-injected mice orally supplemented with either live *F. prausnitzii* and *A. muciniphila* at 2 weeks post-LCWE injection (n=4-10/group). **(D, E)** Spearman or Pearson correlation between the abundance of *F. prausnitzii* and *A. muciniphila* and heart vessel inflation score (D) or maximal abdominal aorta diameter and total abdominal aorta area (Pearson correlation) (E). Data are presented as mean ± SEM and pooled from two independent experiments (A-C). ***$p$<0.001 and ****$p$<0.0001 by unpaired t-test with or without Welch's correction (A), one-way ANOVA with Tukey's multiple comparison tests (B), and Kruskal-Wallis with Dunn's multiple comparisons test (C).

### Figure S5 *(related to Figure 5)*

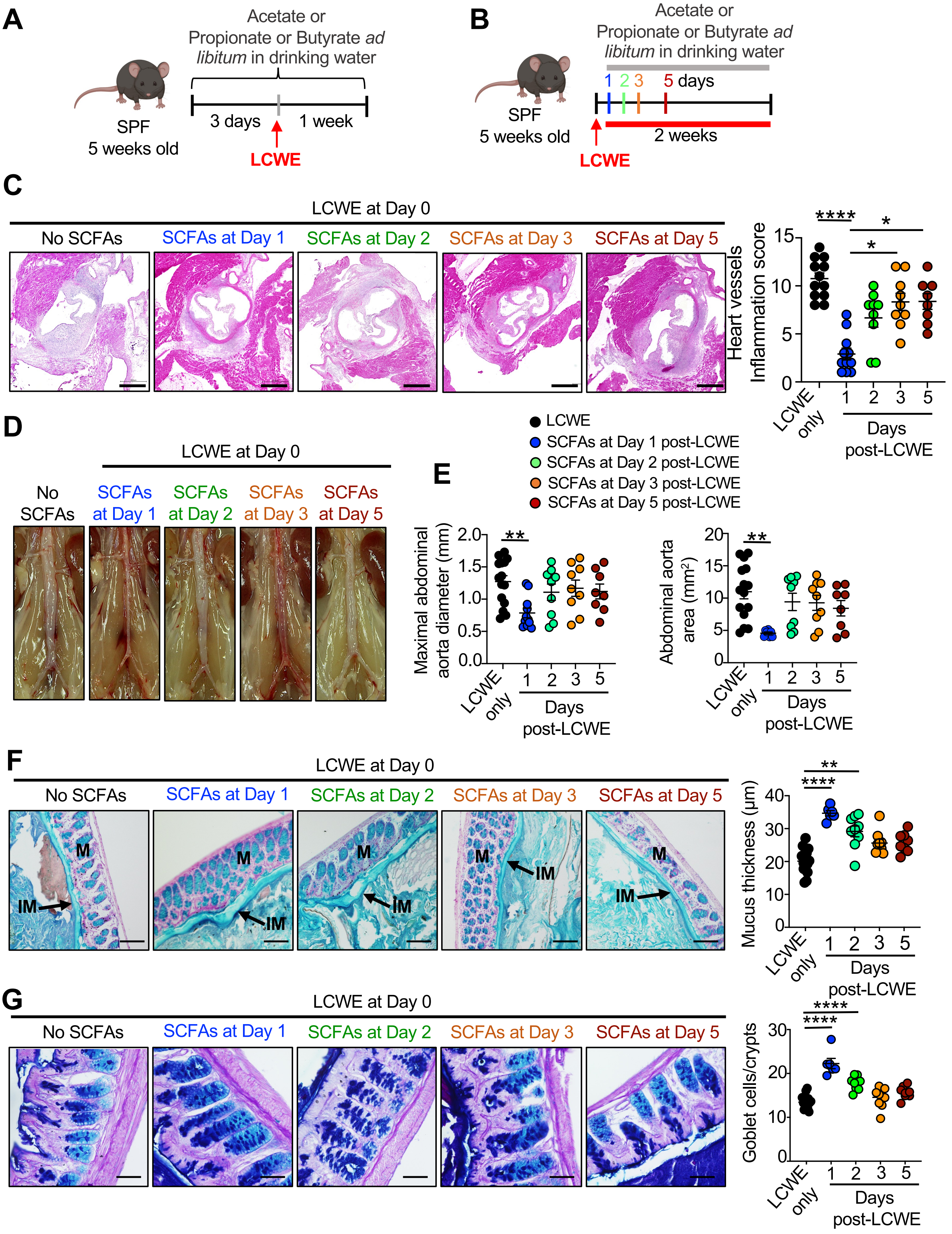

**Supplementary Figure 5 (*related to Figure 5*). Therapeutic SCFAs oral supplementation during LCWE-induced KD vasculitis.**

**(A)** Schematic of the experimental design. Mice received *ad libitum* in drinking water either acetate, propionate, or butyrate starting 3 days before LCWE injection until the experiment endpoint. **(B)** Schematic of the experimental design. Mice received *ad libitum* in drinking water a cocktail of SCFAs composed of acetate, propionate and butyrate starting either on days 1, 2, 3 or 5 after LCWE injection and until the experiment endpoint. **(C)** Representative H&E-stained heart sections and heart vessel inflammation score of LCWE-injected mice and LCWE-injected supplemented with SCFAs starting at either day 1, 2, 3 or 5 post-LCWE injection (n=8-12/group) at two weeks post-LCWE. Scale bars, 500μm. **(D, E)** Representative pictures of the abdominal area (D), maximal abdominal aorta diameter, and abdominal aorta area (E) of LCWE-injected mice and LCWE-injected supplemented with SCFAs starting at either day 1, 2, 3 or 5 post-LCWE injection (n=8-15/group) at two weeks post-LCWE **(F)** Representative photomicrograph showing Alcian blue staining of colon tissues and mucus thickness measurements from colon tissues of LCWE-injected mice and LCWE-injected mice supplemented with SCFAs starting at either day 1, 2, 3 or 5 post-LCWE injection (n=6-15/group) at two weeks post-LCWE. Scale bars; 100μm. **(G)** Representative photomicrograph showing Alcian blue-periodic acid-Schiff (AB-PAS) staining of colon tissues and Goblet cell counts per crypts of LCWE (n=10/group) from tissues from colon tissues of LCWE-injected mice and LCWE-injected supplemented with SCFAs starting at either day 1, 2, 3 or 5 post-LCWE injection (n=8-12/group) at two weeks post-LCWE. Scale bars, 500μm. Data was compiled from two-three independent experiments (C-G) and presented as mean ± SEM. *$p$<0.05, **$p$<0.01, ***$p$<0.001, ****$p$<0.0001 by Kruskal-Wallis with Dunn's multiple comparisons test (C, E, F), and one-way ANOVA with Tukey's multiple comparison test (G). Abbreviation of SCFA, short chain fatty acids.

### Figure S6 *(related to Figures 5 and 6)*

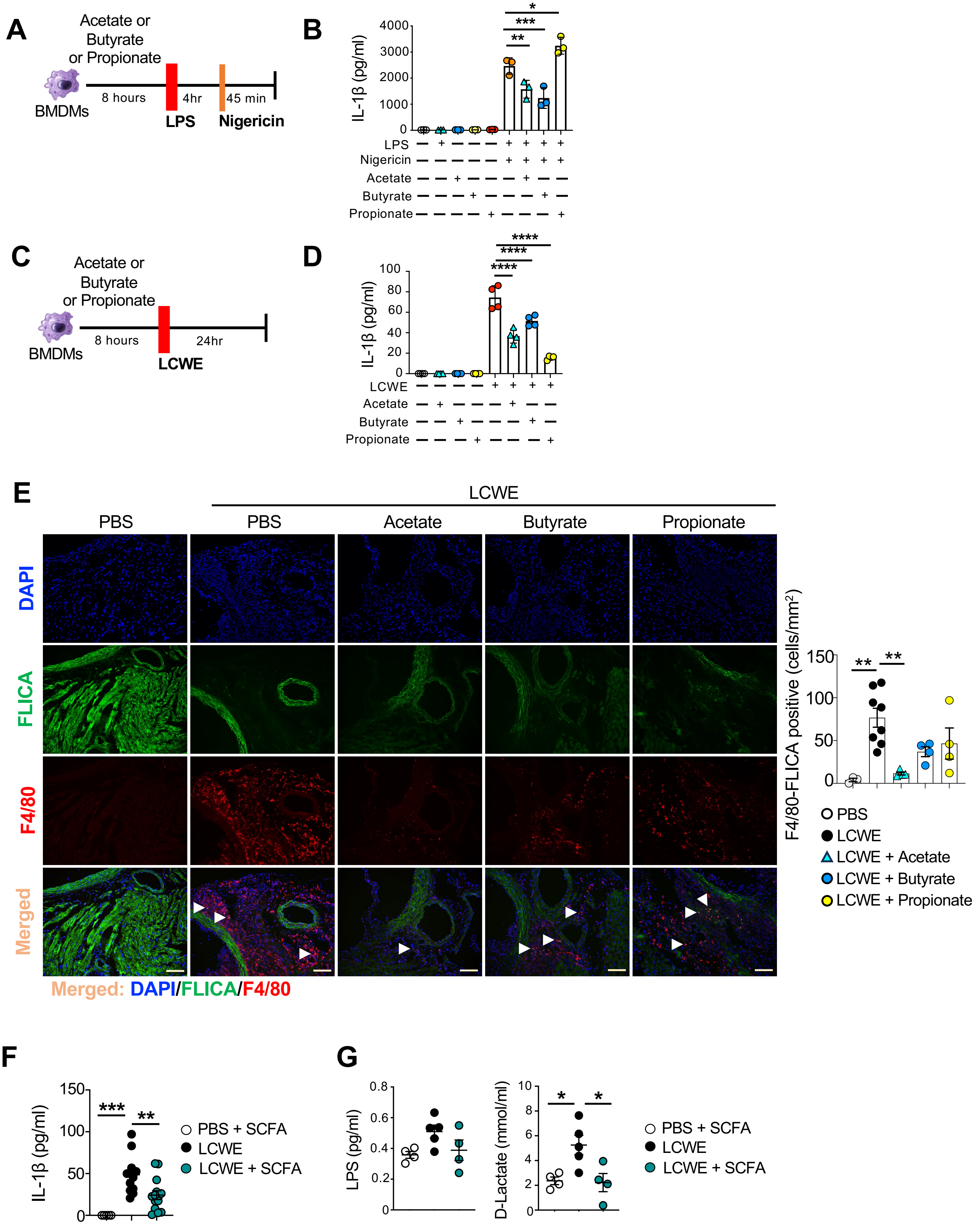

**Supplementary Figure 6 (*related to Figure 5*). SCFAs decrease IL-1β production *in vitro* and *in vivo*. (A)** Schematic of the experimental design. BMDMs were pretreated with either acetate, propionate, or butyrate for 8 hours, primed with LPS, and stimulated with nigericin for 45m minutes, when supernatants were collected, and IL-1β was measured by ELISA. **(B)** Levels of IL-1β produced by BMDMs pretreated with either acetate, propionate, or butyrate, primed with LPS and stimulated with nigericin (n=3/group). **(C)** Schematic of the experimental design. BMDMs were pretreated with either acetate, propionate, or butyrate for 8 hours, and stimulated with LCWE for 24 hours, when supernatants were collected, and IL-1β was measured by ELISA. **(D)** Levels of IL-1β produced by BMDMs pretreated with either acetate, propionate, or butyrate, and stimulated with LCWE (n=4/group). **(E)** Representative picture of F4/80 and FLICA staining and quantification of F480+ FLICA+ cells in heart tissues from either PBS-injected mice, LCWE-injected mice, and LCWE-injected mice that were orally supplemented with either acetate, butyrate, or propionate at one week post-LCWE injection. SCFAs supplementation was started 3 days before LCWE injection. White arrows indicate F4/80+ FLICA+ cells (n =3-8/group). Scale bars: 50μm. **(F)** IL-1β levels in the peritoneal lavage of PBS-injected control mice, LCWE-injected mice, and LCWE-injected mice orally supplemented with a mix of SCFAs (butyrate, acetate, and propionate, provided 3 days before LCWE injection), at 24 hours post-LCWE injection (n=5-14/group). **(G)** Serum levels of LPS and D-Lactate from PBS-injected control mice, LCWE-injected mice, and LCWE-injected mice orally supplemented with a mix of SCFAs (butyrate, acetate, and propionate, provided 3 days before LCWE injection), at 24 hours post-LCWE injection (n=4-5/group). Data was compiled from two independent experiments (B-F) and presented as mean ± SEM. *$p$<0.05, **$p$<0.01, ***$p$<0.001, ****$p$<0.0001 by Kruskal-Wallis with Dunn's multiple comparisons test (C, E) and one-way ANOVA with Tukey's multiple comparisons (F, G).
